# Adaptation across geographic ranges is consistent with strong selection in marginal climates and legacies of range expansion

**DOI:** 10.1101/2020.08.22.262915

**Authors:** Megan Bontrager, Takuji Usui, Julie A. Lee-Yaw, Daniel N. Anstett, Haley A. Branch, Anna L. Hargreaves, Christopher D. Muir, Amy L. Angert

## Abstract

Every species experiences limits to its geographic distribution. Some evolutionary models predict that populations at range edges are less well-adapted to their local environments due to drift, expansion load, or swamping gene flow from the range interior. Alternatively, populations near range edges might be uniquely adapted to marginal environments. In this study, we use a database of transplant studies that quantify performance at broad geographic scales to test how local adaptation, site quality, and population quality change from spatial and climatic range centers towards edges. We find that populations from poleward edges perform relatively poorly, both on average across all sites (15% lower population quality) and when compared to other populations at home (31% relative fitness disadvantage), consistent with these populations harboring high genetic load. Populations from equatorial edges also perform poorly on average (18% lower population quality) but, in contrast, outperform foreign populations (16% relative fitness advantage), suggesting that populations from equatorial edges have strongly adapted to unique environments. Finally, we find that populations from sites that are thermally extreme relative to the species’ niche demonstrate strong local adaptation, regardless of their geographic position. Our findings indicate that both nonadaptive processes and adaptive evolution contribute to variation in adaptation across species’ ranges.

## Introduction

Geographic ranges form as populations of a species spread across a landscape, via successive dispersal and adaptation to local environments. Capacity for rapid range expansion is well documented, for example, when species colonize a new continent after human introduction (Stuckey, 1980; Lachmuth et al., 2010), or recolonize formerly glaciated areas (Clark, 1998, and references within). Studies of population differentiation along geographic gradients provide classic examples of adaptation to variation in the environment (Turesson et al., 1930; Clausen et al., 1948). The extent to which range expansions require adaptation to new environments or simply dispersal into already suitable environments remains an open question (Sheth et al., 2020). Regardless, whether adaptation is necessary for expansion or honed after colonization, spread and adaptation do not happen indefinitely. This implies that the balance of ecological and evolutionary forces shifts across species’ ranges, ultimately preventing further adaptation and halting spread.

Several hypotheses have been proposed for why adaptation might stall at niche-limited range edges. These non-mutually exclusive hypotheses are based on population genetic and demographic factors, and each predicts a failure in local adaptation at the edge. Most assume that species’ ranges overlay an ecological gradient (such as climate or the abundance of competitors) that results in smaller populations toward range edges (Brown, 1984; Gaston, 2009). Small populations are more vulnerable to genetic drift and demographic stochasticity (Eckert et al., 2008; Kawecki, 2008). Edge populations could also be pulled away from their local fitness optima if net migration occurs mostly from central towards edge populations (known as ‘gene swamping’, Kirkpatrick and Barton, 1997). The combination of drift and gene swamping may limit adaptation and lead to a temporally stable range edge, especially when underlying environmental gradients are steep relative to dispersal distances (Polechová and Barton, 2015; Polechová, 2018, see Bachmann et al. 2020 for an empirical test). Furthermore, if range-edge sites are uniquely challenging, adaptation might also be hampered by reduced genetic variation following strong and persistent directional selection (Hoffmann and Blows, 1994), or by genetic constraints arising from antagonistic genetic correlations (Levins, 1968; Antonovics, 1976; Bradshaw and McNeilly, 1991, see Olsen et al. 2019 for an empirical test). Together, this body of theory predicts that the magnitude of local adaptation decreases from geographically central to peripheral populations (Fig. 1A). The patterns and processes invoked by these equilibrial hypotheses have mixed support (Bridle et al., 2009; Pironon et al., 2017) and estimates of how local adaptation varies spatially across geographic ranges (e.g., Putnam and Reich, 2017, Fig. 2A) remain rare.

**Fig. 1.**
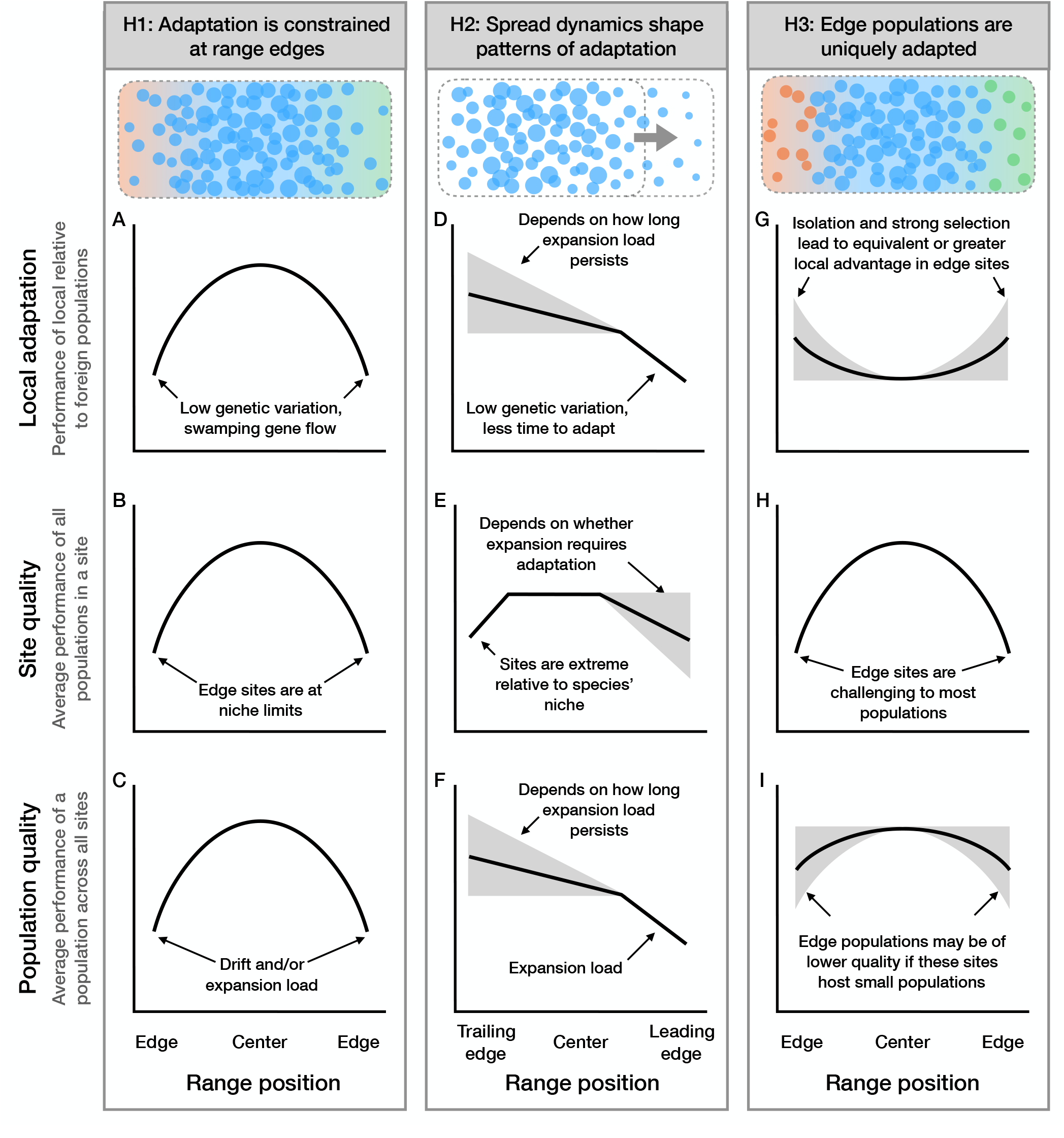
Predicted relationships between geographic range position and local adaptation, site quality, or population quality derived from three hypotheses about evolutionary processes across ranges. Hypothesis 1 describes a scenario where range edges are constrained by adaptation, predicting: (A) declines in local adaptation towards range edges (regardless of whether edges are latitudinal, elevational, or geographic), (B) declines in site quality at range edges because they are ecologically challenging (i.e., marginal relative to the species’ niche), and (C) declines in population quality towards range edges because small edge populations accumulate genetic load through inbreeding or drift. Hypothesis 2 describes ranges that have undergone expansion in one direction, predicting: (D) declines in local adaptation towards high latitudes and elevations (leading edges) but not towards non-directional peripheries (which pool rear and leading edges together), (E) decines in site quality at leading edges (if novel adaptation is required for spread) and/or at rear edges (if environmental change is shifting these sites outside the species’ niche), (F) lower population quality at leading edges due to genetic load, particularly expansion load. Hypothesis 3 describes a scenario where edge populations are uniquely adapted to marginal environments, predicting: (G) equivalent or even increasing local adaptation towards geographic peripheries (regardless of direction), latitudinal edges, and elevational edges, (H) declines in site quality at range edges because they are ecologically challenging (i.e., marginal relative to the species’ niche), and (I) equivalent (if selection has not reduced population sizes) or decreasing (if selection has reduced population sizes) population quality at range edges. Circles (top row) represent populations in a cartoon range (dashed rectangle), where circle size indicates population size and the color gradient indicates a spatially varying environment to which edge populations are maladapted (H1) or strongly adapted (H3). Shaded areas in D and F depict uncertainty in how long expansion load persists after the expansion front passes through (Peischl et al., 2013).

**Fig. 2.**
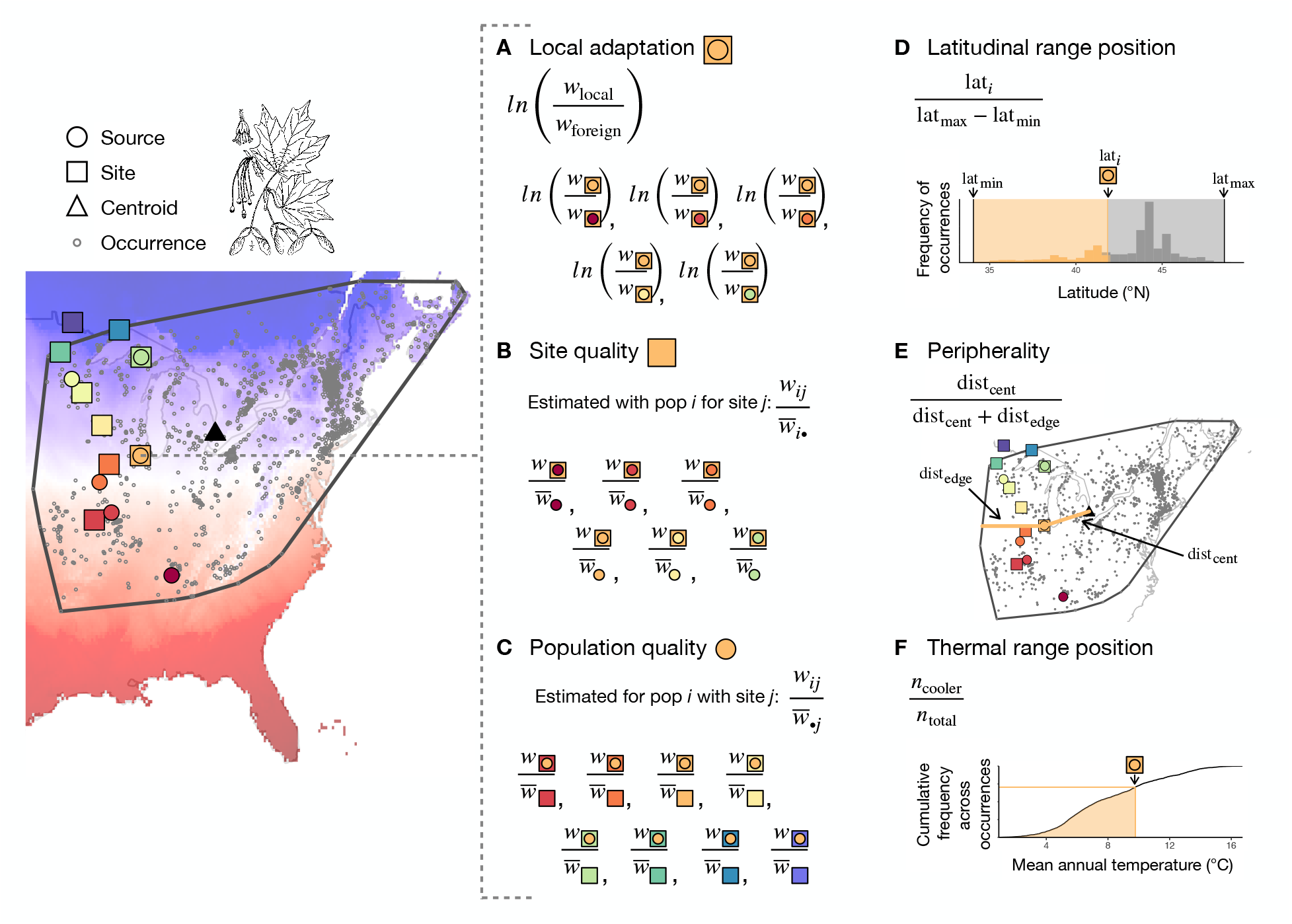
Schematic of methods for calculating response variables (A-C) and predictor variables (D-F), illustrated for data from sugar maple, *Acer saccharum*. Transplant sources (circles) and sites (squares) from Putnam and Reich (2017) are overlain on records of occurrence (grey dots) from GBIF (www.gbif.org) and mean annual temperature (red-blue color gradient) from Worldclim (Fick and Hijmans, 2017). For each transplant site, (A) local adaptation is estimated as the fitness advantage of the local source population over each foreign source and (B) site quality is estimated as the mean fitness of all sources transplanted to that site, after normalizing each source’s performance to its experiment-wide average. For each source population, (C) population quality is estimated as the mean fitness of that population across all sites where it was transplanted, after first normalizing each population’s site-specific performance to the mean fitness of all sources at that site. For each source and site, locality records are used to estimate the (D) latitudinal range position based on their position relative to the maximum and minimum values observed across all localities and (E) peripherality relative to the centroid (triangle) and edge of the minimum convex polygon (grey line) surrounding all occurrences. Climate values associated with each occurrence record were used to calculate the cumulative frequency of mean annual temperature values across the species’ range, with (F) thermal range position estimated based on its rank within the frequency distribution. Elevation was handled analogously to latitude, and mean annual precipitation analogously to temperature.

The hypotheses above predict that the selective challenge posed by sites will increase toward range edges (lower ‘habitat quality’ *sensu* Blanquart et al., 2013, hereafter ‘site quality’), while the genetic quality of populations (‘deme quality’ *sensu* Blanquart et al., 2013, hereafter ‘population quality’) will decrease. In this context, site quality is a property of the site relative to the species’ niche. Empirically, the relative quality of sites can be approximated based on the average performance of multiple populations transplanted to those sites (Fig. 2B). Sites that are marginal relative to a species’ niche should pose a greater fitness challenge for individuals, even after accounting for the effects of local adaptation and population quality (i.e., across multiple populations of a given species, average fitness will be low in a low-quality site). It follows that if low-quality sites predominate at range edges (Fig. 1B), selection at edges will be stronger than in the range center (Crow, 1958; Caruso et al., 2017; Angert et al., 2020). However, if response to selection is constrained, individual fitness and population size will remain low in these low-quality sites. The resulting gradient in population size could foster genetic drift or swamping gene flow, which could further exacerbate the size gradient. As a result, low site quality at range edges should reduce population quality, as small populations will experience stronger drift (Koski et al., 2019, Fig. 1C). Empirically, the relative quality of populations can be approximated based on the average performance of each population transplanted to multiple sites (Fig. 2C).

The theory discussed so far is focused on explaining stable (equilibrial) range limits. However, some range edges are not at equilibrium and instead reflect ongoing range shifts, such as those driven by glacial retreats and climate change. The legacy of current or past range shifts should leave signatures on geographic patterns of adaptation and population quality. In theory, serial founder events at the leading edge of a recent range expansion can lead to allele surfing (Excoffier et al., 2009) and the fixation of deleterious alleles (i.e., expansion load, Peischl et al., 2013), thus reducing population quality (Fig. 1F). Genomic signatures of expansion load have been documented in several systems (Zhang et al., 2016; González-Martínez et al., 2017; Perrier et al., 2020), and have also been tied to reduced fitness in at least one system (*Arabidopsis lyrata*, Willi et al., 2018). In cases where niche evolution is required during range expansion, site quality will be low at leading edges (Fig. 1E). Serial founder events during expansions can reduce genetic variation and thereby decrease evolutionary potential for local adaptation (Excoffier et al., 2009, Fig. 1D). For example, post-Pleistocene range expansion is hypothesized to explain the reduced additive genetic variation and lower response to selection displayed in Spanish and Portuguese populations of annual mercury (*Mercurialis annua*) following colonization of the Iberian peninsula (Pujol and Pannell, 2008). Along environmental gradients, leading-edge populations may also exhibit low local adaptation because they have had less time to adapt to their new environment.

Different processes might be expected to operate at rear edges, where sites may be extreme (or becoming extreme) relative to the species’ niche. Here conditions may be challenging to all populations (low site quality; Fig. 1E). However, in contrast to expectations at leading edges, rear-edge populations may be best adapted to local conditions (Fig. 1D), having had longer to adapt and perhaps possessing phenotypes that are already best suited to the direction of environmental change (relative to populations elsewhere). Agnostic to the type of edge, similar patterns have been hypothesized for edge populations in general if they occupy sites that are environmentally distinct relative to others across the range (Fig. 1H) and have unique adaptations that allow them to persist there (Rehm et al., 2015; Roschanski et al., 2016, Fig. 1G)—an idea that has been proposed to increase the conservation value of edge populations (Lesica and Allendorf, 1995; Gibson et al., 2009). These populations may be of no lower quality than populations elsewhere in the range if strong selection did not cause population size contractions (Fig. 1I), or population quality may be low at edges if challenging sites host smaller populations with increased genetic load (Fig. 1F, I). Overall, these predictions suggest that populations from some range edges may be uniquely adapted to challenging or climatically marginal sites, but that occupying such sites may reduce population quality.

Some of the aforementioned mechanisms shaping adaptation across geographic ranges operate based on the dynamics of spatial spread alone (e.g., expansion load). Others are based on the response—or lack thereof—to divergent selection at range edges, and assume that geographic peripherality is equivalent to ecological marginality. Across large spatial scales, climate is often assumed to be the major ecological agent of divergent selection and a predominant factor limiting species’ ranges (Araújo and Rozenfeld, 2014; Cunningham et al., 2016). However, climate marginality is only one component of ecological marginality, and increasing evidence implicates biotic interactions as a key driver of selection even at broad geographic scales (Araújo and Rozenfeld, 2014; Belmaker et al., 2015; Armitage and Jones, 2020; Angert et al., 2020, and references therein). Additionally, it is not always the case that geographically peripheral populations inhabit climatically marginal conditions (Lira-Noriega and Manthey, 2014; Pironon et al., 2015; Oldfather et al., 2020). Theory is only beginning to extend to situations where geographic and ecological marginality are decoupled (e.g., Polechová, 2018), and it remains an open question whether variation in population quality and local adaptation are better predicted by ecological or spatial marginality. For example, when response to selection is not limited by genetic constraints that arise from spatial processes (such as gene flow or expansion) and climate is the major selective force across large spatial scales, local adaptation should be more strongly predicted by climate than by spatial proxies. In contrast, when expansion dynamics predominate as a species colonizes climatically suitable areas, local adaptation should be more strongly correlated with spatial than environmental predictors.

We have synthesized models into three competing hypotheses that make contrasting predictions about how local adaptation, site quality, and population quality vary across the range (Fig. 1). To test these competing hypotheses, we leverage collection records and published transplant studies. Many studies have used transplants among natural populations to investigate local adaptation (Hereford, 2009). These studies use the fitness of a local population relative to that of foreign populations as a quantitative metric for how well-adapted that population is to its local environment (Kawecki and Ebert, 2004, Fig. 2A). Site quality (Fig. 2B) and population quality (Fig. 2C) can also be estimated from transplant data and provide complementary insights into evolution across spatial and environmental gradients (Blanquart et al., 2013, Fig. 1). For each transplant study, we use collection records to estimate the position of source populations and transplant sites with respect to species’ geographic and climatic range (Fig. 2D-F). We then compare these range position estimates to transplant fitness to test three hypotheses about equilibrial and non-equilibrial ranges and the interplay of ecological marginality and evolutionary processes across species’ distributions:

**Hypothesis 1**: Adaptation is constrained at range edges. This hypothesis predicts that local adaptation, site quality, and population quality are lower at range edges because swamping gene flow or lack of appropriate genetic variation constrain adaptation and result in niche-limited ranges. From this hypothesis, we derive the testable predictions illustrated in Fig. 1A-C).

**Hypothesis 2**: Spread dynamics shape patterns of adaptation across ranges that are not at equilibrium. Local adaptation is weaker at leading edges because they are younger or have lower genetic variation, but stronger at rear edges because they have had more time to respond to selection or because shifting conditions favor individuals already partially adapted to extreme conditions. Predictions stemming from this hypothesis are illustrated in Fig. 1D-F).

**Hypothesis 3**: Edge populations are uniquely adapted. Local adaptation is equal across ranges or even higher at range edges because isolation and strong selection in peripheral, low-quality sites promote divergence. Specific predictions from this hypothesis are illustrated in Fig. 1G-I).

We also test whether climatic marginality (i.e., climates that are in the tails of the distribution of occupied climates across a species’ range) is a major driver of large-scale spatial differences in local adaptation, site quality, and population quality. The comparison of climatic versus spatial predictors can reveal the importance of non-climatic forms of environmental marginality at spatial edges as well as environment-independent drivers, such as population age and expansion load. Finally, we examine correlations between spatial and climatic marginality across the ranges of species in our dataset.

## Methods

### Literature search

We used a database of studies that had conducted transplant experiments moving populations of terrestrial species at geographic scales, which was previously compiled to examine adaptation to biotic interactions (Hargreaves et al., 2020) and to temporal variation in climate conditions (Bontrager et al., 2020). We searched for studies through 19 March 2017 that moved at least one population to multiple locations or multiple populations to at least one location and subsequently measured at least one component of fitness (see below). Details of literature search methods and criteria for inclusion are in Hargreaves et al. (2020), except that here we excluded 20 studies for which (i) we were unable to resolve taxonomy well enough to confidently gather occurrence information, (ii) inadequate occurrence records were available, or (iii) the study locations encompassed *<*10% of the species’ geographic or elevational range (based on comparisons to independent range maps or descriptions). This resulted in a final dataset of 119 papers representing 135 species (Table S1).

### Data collection

#### Fitness data

Within a study, we collected fitness data for each unique combination of species, site, source, and fitness metric. A ‘site’ refers to the test location, and a ‘source’ refers to the home location of a population. We included only sites and sources that were within a species’ native range (using the characterizations of the original authors when locations were near range edges). When multiple fitness metrics were presented, we collected data from one representative measure in each of five categories (germination, establishment, survival, reproduction, or a composite fitness metric that incorporates at least survival and reproduction, such as population growth rate or lifetime fitness). We used all available metrics from a study in our analyses (see Statistical analyses). Details of fitness data collection can be found in the Supplementary Methods.

#### Source and site geographic data

We gathered location data (latitude, longitude, and elevation) for each source and site. In cases where only the latitude and longitude were provided by authors, we used Google Earth and landmarks to estimate elevation. If latitudes and longitudes were not provided (neither in the paper nor after emails to authors) we estimated locations from maps or site descriptions if possible.

#### Range data

Locality data to define the range of each species were downloaded from the Global Biodiversity Information Facility (GBIF; www.gbif.org) on 1-3 March 2018; digital object identifiers for these data are listed in Table S1. We cleaned these data to remove erroneous localities (e.g., in oceans) and filtered them to each species’ native range. Details of locality cleaning can be found in the Supplementary Methods. After cleaning, the number of localities retained for each species ranged from 14 to 35969 (median = 1499).

#### Climate data

We extracted climatic data for all sources, sites, and localities of every species from Bioclim (version 2.1, 30 second resolution, Fick and Hijmans, 2017). We used 1960-1990 averages of mean annual temperature (BIO1, hereafter ‘MAT’) and mean annual precipitation (BIO12, hereafter ‘MAP’).

### Response variables

#### Local adaptation

We calculated local adaptation as the log-transformed ratio of the local population’s fitness (*w*_local_) to the fitness of each foreign population (*w*_foreign_) transplanted to a site, i.e., the magnitude of local advantage (Fig. 2A). Local adaptation was calculated with each fitness metric available for each foreign population at each site, yielding a total of 3723 data points. Positive values of this metric indicate local adaptation, while negative values indicate foreign advantage. When the average fitness of either population is very low, it will often be measured as 0 due to finite sample sizes. This results in an estimate of local adaptation that is non-finite. Therefore, we treated these observations as censored (implementation of censoring in Stan is described in section 4.3 of the Stan User’s Guide, Stan Development Team, 2018). We used the lowest observed non-zero fitness of any population at a site (*w*_min_) as an estimate of the lowest observable fitness, and calculated censors for our local adaptation metric using this minimum in the ratio. When a foreign population’s fitness was measured as 0 (3.6% of data points), we right-censored the data beyond the value of log(*w*_local_*/w*_min_). When the local population’s fitness was measured as 0 (1.9% of data points), we left-censored the data below values of log(*w*_min_*/w*_foreign_). We then used the Student *t* cumulative density distribution to estimate the likelihood of these observations as the cumulative probability between ∞ and the right-censor (when *w*_foreign_ = 0) or -∞ and the left-censor (when *w*_local_ = 0).

#### Site quality

We estimated site quality based on how populations performed at a given site relative to their average performance across all sites (Fig. 2B). To calculate relative performance, we divided the fitness of a given population at a site by that population’s average fitness across all sites in which it was tested, excluding its home site. We excluded home sites from the denominator because their inclusion tends to bias downward our estimates of population quality for populations tested at few sites, since populations tend to do well in their home sites. Populations that had fitnesses of 0 across all ‘away’ sites were excluded from this analysis (0.8% of observations). This resulted in 7014 observations of relative fitness across 406 sites. We then log-transformed these values so that residual variation would be normally distributed. As a result, sites where there is positive relative performance are those where most populations perform better than their experiment-wide average. Sites where there is negative relative performance are those where populations perform worse than their average. As in calculations of local adaptation, when fitness in a given site was measured as 0 (6.1% of observations), this resulted in a non-finite value (-∞) once transformed. We censored these observations as described for our metric of local adaptation.

#### Population quality

Population quality was estimated based on how a given population performed relative to other populations in the sites where it was tested (Fig. 2C). To calculate relative performance, we divided the fitness of a given population at a site by the average fitness of all foreign populations tested at that same site. We included only foreign populations in the denominator because the inclusion of local populations tends to bias downward our estimates of site quality for sites at which few populations were tested, since populations local to a site tend to do well. Test sites where all foreign populations had fitness of 0 were excluded from this analysis (1.6% of observations). This resulted in 7097 observations of relative fitness across 1510 populations. We then log-transformed these values so that residual variation would be more normally distributed. Populations that have positive relative performance are those that frequently outperform others across test sites. As with other response metrics, when fitness of a focal population was measured as 0 (4.7% of observations), this resulted in a non-finite value (-∞) once transformed. We censored these observations as described above.

### Predictor variables

#### Spatial range position

We estimated three categories of spatial range position: latitudinal range position, elevational range position, and geographic peripherality. We calculated these for each transplant site and each source population.

We calculated a metric of latitudinal range position relative to the maximum and minimum latitudes of the species’ localities (Fig. 2D). Latitudinal position is bounded by 0 and 1, with 0 being the equatorial limit, and 1 being the poleward limit. For the four taxa in our dataset with ranges spanning the equator, we designated the limit further from the equator as the poleward limit. Transplant sites or sources beyond the equatorial or poleward latitudinal limits encompassed by the species’ localities were assigned 0 or 1, respectively.

We calculated the elevational range position of sites and sources analogously. We primarily used maximum and minimum elevations from the literature because many species had few locality records that reported elevation. However, we used GBIF records when they expanded the published range or when published elevational ranges were unavailable. Locations with an elevational range position value of 0 are at the lower limit of the species’ elevational range and those with a value of 1 are at the upper limit. We did not calculate elevational ranks for the 15 species in our dataset that had ten or fewer records with elevations. The species for which we did calculate elevational ranks had between 11 and 13808 records with elevation data (median = 231).

We defined geographic peripherality as the position of each transplant site relative to the center and the edge of the focal species’ range (Fig. 2E). Specifically, we calculated the distance from the transplant site to the centroid of the minimum convex polygon (MCP) encompassing all locality points for the species in question. We also calculated the minimum distance from the transplant site to the edge of the MCP. Peripherality was then calculated as the distance to the centroid, divided by the sum of the distance to the centroid and the distance to the edge (following Griffin and Willi, 2014; Lee-Yaw et al., 2017). Peripherality ranged from 0 to 1, with larger values indicating greater proximity to the range edge. MCPs for each species were generated using the mcp function from the adehabitatHR package version 0.4.15 (Calenge, 2013) in R. Measures of distance were made following re-projection of the locality and transplant site data to a region-appropriate equal area projection.

#### Climatic position

We characterized the climatic position of each transplant site and source by their ranks in the empirical cumulative distributions of MAT and MAP (Bioclim variables, Fick and Hijmans, 2017) observed across all the locality records for a given species (Fig. 2F). Climatic position is bounded by 0 and 1, with 0 being the coldest or driest recorded site and 1 being the warmest or wettest recorded site.

#### Covariates

We calculated several covariates that are likely to influence local adaptation or our metrics of site or population quality. We included the absolute value of the latitude of a site or source as a proxy for the potential influence of post-glacial dynamics (Svenning and Skov, 2004; Muellner et al., 2005; Souto et al., 2015). The magnitude of local adaptation and differences in quality detected in transplant studies may be correlated with the magnitude of environmental differences between the populations selected for the study (Hereford, 2009). To account for this, we calculated the geographic distance, absolute temperature difference, and absolute precipitation difference between each source and site. For models of local adaptation, we also included the temperature deviation between a site’s historical conditions and those during the experiment, which could alter the strength of local advantage (calculation methods are described in Bontrager et al., 2020).

### Statistical analyses

All statistical analyses were conducted in R version 3.6.2 (R Core Team, 2019). We first calculated Pearson correlations amongst all predictor variables to (a) assess possible collinearity and (b) test whether spatial peripherality is associated with climatic marginality.

Given low correlations amongst predictors (see Results), we were able to all include all predictors in our models. To test how local adaptation varies with range position (Fig. 1A, D, G), we included as predictors three metrics of spatial position (latitudinal range position, elevational range position, peripherality), two metrics of climatic position (temperature range position, precipitation range position), and five covariates (geographic distance, temperature difference, precipitation difference, absolute value of latitude, and temperature deviation). Spatial and climatic predictors (as well as the covariates of the absolute value of latitude and the temperature deviation) describe the local population (i.e., the transplant site). The covariates of geographic distance, temperature difference, precipitation difference describe the foreign population (relative to the transplant site) in a given comparison. We fit quadratic effects of latitudinal, elevational, temperature, and precipitation positions because low and high values of these variables both represent range edges. For geographic peripherality, we fit a linear term only because low values represent the range center and high values represent the range edge. We included only linear effects of the covariates absolute latitude, geographic distance, and temperature and precipitation differences because we predicted linear changes towards higher latitudes and with greater distances. All continuous variables were centered to a mean of zero and scaled to a standard deviation of one, and were back-transformed to their original scale for plotting. We modeled responses using a Student *t* distribution, a robust regression approach which is less sensitive to extreme data points (Gelman et al., 2013), which are present in the distributions of our response variables. We also included a group level intercept for each combination of study, site (or source when the response was population quality), species, and fitness metric to account for non-independence within these groups. Linear mixed-effect regression models were fit with Stan version 2.19 (Stan Development Team, 2018) and were run for 20000 iterations until all parameters had converged 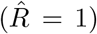. Trace plots were visually inspected and posterior prediction plots were checked for consistency with the data (Gelman et al., 2013). Data and code are available upon request during review and will be deposited in a public data archive upon publication.

Models for site quality (Fig. 1B, E, H) and population quality (Fig. 1C, F, I) were analogous, with the following exceptions. First, for population quality, spatial and climatic positions (as well as absolute latitude) refer to where the population originates, whereas for local adaptation and site quality, values are based on the positions of the transplant site. Second, we only included temperature anomaly as a covariate in models of local adaptation. Finally, for population quality, we may only detect an effect of a given range axis when that range axis is well-sampled within a transplant site. Similarly, we may only be able to detect effects of range axes on site quality when we estimate it from populations transplanted to sites that span a large portion of a given range axis. So, for each predictor of interest, we filtered the dataset to observations falling above the 70th percentile of range coverage on that range axis. This threshold was chosen to balance maintaining a large sample size (∼2000 lines of data) with using the best data to test each hypothesis, and resulted in a different subset of the data and a different model fit for each combination of predictor and response. We included all predictors in these models, but focus our interpretation on the predictor that has good coverage in each dataset (i.e., the predictor for which we filtered the data).

### Sensitivity tests

We examined the sensitivity of our results (i.e., we observed whether they changed qualitatively) to various combinations of alternative range position metrics. First, although we favored an MCP around occurrence records to characterize peripherality because it requires no parameters to be set by the user, it does not necessarily preserve range shape and might encompass large areas where a focal species is absent. We conducted a sensitivity test in which we calculated peripherality based on the intersection between the species’ MCP and a polygon representing buffered locality data points (Supplementary Methods, sensitivity test 1). Second, latitudinal and elevational positions based on maximum and minimum observations could be sensitive to outlier localities (i.e., disjunct populations beyond the species’ main geographic range), so we conducted sensitivity tests with rank-based metrics. Latitudinal (or elevational) rank was defined as the rank of a given site or source in the empirical cumulative distribution of latitudes (or elevations) from the set of localities for a species (i.e., rank transformed; sensitivity test 2). Though ranks should be less sensitive to outliers, they are likely to be influenced by spatially uneven sampling and reporting of locality data across species’ ranges. Finally, we used an alternative metric of climatic position based on temperature and precipitation maximums and minimums (sensitivity test 3). However, we favored climatic ranks because they capture the prevalence of different climatic conditions across a species’ range.

## Results

### Properties of the final dataset

The final dataset encompassed 7544 records from 1524 unique source populations moved among 421 sites. On average, a source population was moved to 4 sites (median: 4, range: 1 to 16) while 9 populations were moved to a given site (median: 4, range: 1 to 241). Transplants encompassed up to 15^*°*^C difference in mean annual temperature and 2580 mm difference in mean annual precipitation (Table S2B). Studies varied in the fraction of the species’ range covered by sources and sites, with medians (means) of 20% (31%) for latitudinal ranges, 25% (35%) for elevation ranges, 52% (52%) for precipitation ranges, and 36% (44%) for thermal ranges. Different types of range edges also differed in how frequently they were represented among sites and sources. Using a threshold that defined an edge site as within 10% of the maximum (or minimum) for a given range position metric, 14 (19)% of studies included at least one site (source) near the high latitude edge, 10 (14)% near the low latitude edge, 14 (21)% near the high elevation edge, 48 (56)% near the low elevation edge, 41 (51)% near non-directional peripheries, 24 (31)% near the warmest occurrence, 36 (46)% near the coolest occurrence, 31 (40)% near the wettest occurrence, and 28 (41)% near the driest occurrence. Although six continents were represented in the dataset, study regions were biased towards the northern hemisphere (Table S2D). The 135 study species were heavily skewed towards plants (Table S2E). Approximately 40% of the studies yielded a lifetime or composite estimate of fitness encompassing both survival and reproduction (Table S2F). When filtered to the subsets used for population and site quality analyses, between 1447 and 2119 observations were retained in each subset.

### Local adaptation is lower at poleward edges but higher in thermally extreme sites

Consistent with Hypothesis 2, local adaptation declined towards poleward latitudinal and upper elevational range edges but was unrelated to geographic peripherality (Fig. 3A-C, Table S3). On average, local populations showed a 1.10 to 1.16-fold fitness advantage at equatorward edges and central sites but foreign populations had a 1.44-fold advantage at poleward edges (Fig. 3A). Populations also showed higher local advantage at low elevation edges than central sites (from 1.34-fold local advantage at low edges to no local advantage at elevation midpoints, with high uncertainty in estimates at upper edges; Fig. 3B). In thermally marginal sites, the magnitude of local advantage was 20% (warm edges) to 45% (cool edges) greater than in thermally central sites (where local populations had a 10% fitness advantage; Fig. 4A, Table S3). This result parallels Hypothesis 3, but at thermal rather than spatial edges. Local adaptation was unrelated to precipitation position (Fig. 4B, Table S3). Effects of other covariates are reported in the supplementary results and Table S3. Results for local adaptation were robust to most sensitivity tests (see supplementary results, Table S14, Fig. S1, S2).

**Fig. 3.**
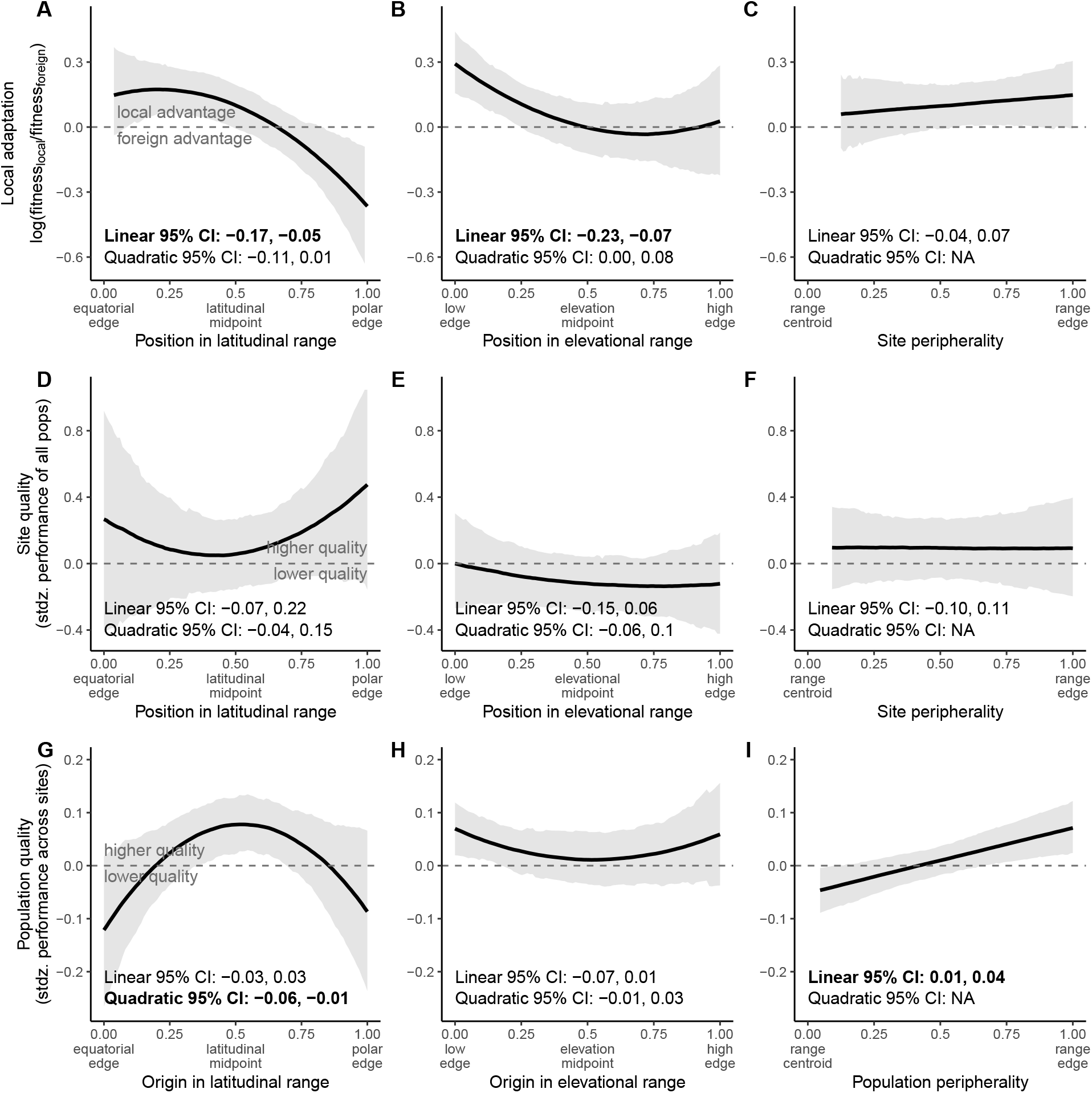
Local adaptation (A-C), site quality (D-F), and population quality (G-I) across dimensions of geographic range position defined by latitude (A, D, G), elevation (B, E, H), or a directionless peripherality metric (C, F, I). Solid black lines and grey shading depict median and 95% highest density posterior interval, respectively, evaluated at mean values of the covariates. Bold font indicates 95% intervals that do not encompass 0 for linear or quadratic terms of the predictor variable. The plotted regression lines result from equations including both linear and quadratic terms (when applicable), regardless of whether estimates for these terms differ from 0. Dashed horizontal line at y=0 depicts the threshold above which populations are more fit, on average, than foreign populations at a site (A-C), than all populations in an average site (D-F), or than the average population across all sites (G-I).

**Fig. 4.**
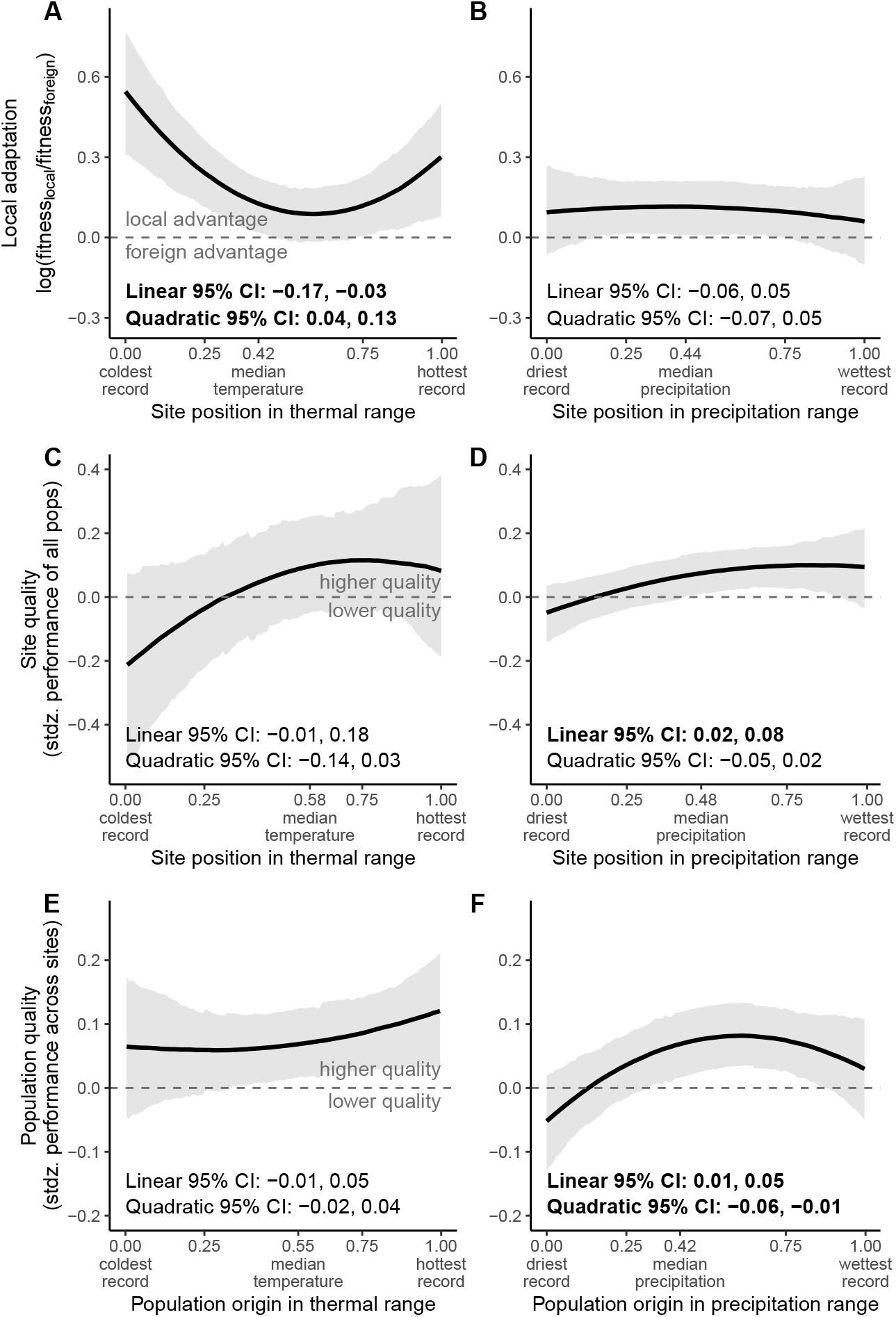
Local adaptation (A, B), site quality (C, D) and population quality (E, F) across dimensions of climatic range position defined by mean annual temperature (A, C, E) or a mean annual precipitation (B, D, F). Solid black lines and grey shading depict median and 95% highest density posterior interval, respectively, evaluated at mean values of the covariates. Bold font indicates 95% intervals that do not encompass 0 for linear or quadratic terms of the predictor variable. The plotted regression lines result from equations including both linear and quadratic terms (when applicable), regardless of whether estimates for these terms differ from 0. Dashed horizontal line at y=0 depicts the threshold above which (A, B) the local population is more fit, on average, than foreign populations, or (D, E) mean fitness is greater than the average site, or (E, F) mean fitness is greater than the average population.

### Site quality is lower in some marginal climates but not at spatial range edges

Site quality did not vary with spatial range position (Fig. 3D-F, Tables S4, S5, S6). Site quality tended to decline (by 28% on average) in climates that are cool relative to the center of species’ temperature niches, though there was large uncertainty in this parameter estimate, and the credible interval just included the possibility of no effect (Fig. 4C, Table S7). Site quality declined by about 13% in climates that are dry relative to the center of species’ precipitation niches (Fig. 4D, Table S8; similar to Hypothesis 3, but with low site quality at climatic rather than spatial edges). Using an alternative calculation of climatic position (with sites characterized relative to maximum and minimum climate values, rather than their rank in a distribution of localities), all climatic edges had lower site quality (supplementary results, Table S14, Fig. S2C, D). Effects of other covariates are also reported in the supplementary results.

### Population quality is lower at latitudinal, upper, warm, and wet edges

Population quality peaked at latitudinal range centers and on average declined by 18% towards equatorward edges (consistent with Hypotheses 1 and 3) and by 15% towards poleward edges (Fig. 3G, Table S9). The significance of this pattern was sensitive to how range position metrics were calculated, but the trend was consistent across tests (Table S14, Fig. S1G). Population quality increased towards non-directional peripheries (Fig. 3I, Table S11) and declined towards the extremes of species’ precipitation niches, particularly dry edges (Fig. 4F, Table S13), though these patterns were sometimes sensitive to alternative metrics of range position (supplementary results, Table S14, Fig. S1I, S2F). Effects of other covariates are reported in the supplementary results.

### Spatial range edges are not necessarily climatically marginal

Correlation coefficients between spatial position and climatic position were low to moderate (Fig. 5), indicating that range edge populations do not necessarily reside in climatically marginal sites. Equatorward and low-elevation edges tended to be warmer, on average, than most sites where a species occurs, while poleward and high-elevation edges tended to be cooler on average (Fig. 5A, C). However, many interior sites were thermally marginal, and many edge sites were not markedly warm or cold relative to other places where the species occurs (Fig. 5A, C). Correlations between spatial position and precipitation marginality were even lower (Fig. 5B, D). Peripherality, without regard to a latitude or elevation axis, was not associated with climatic marginality (Fig. 5E, F).

**Fig. 5.**
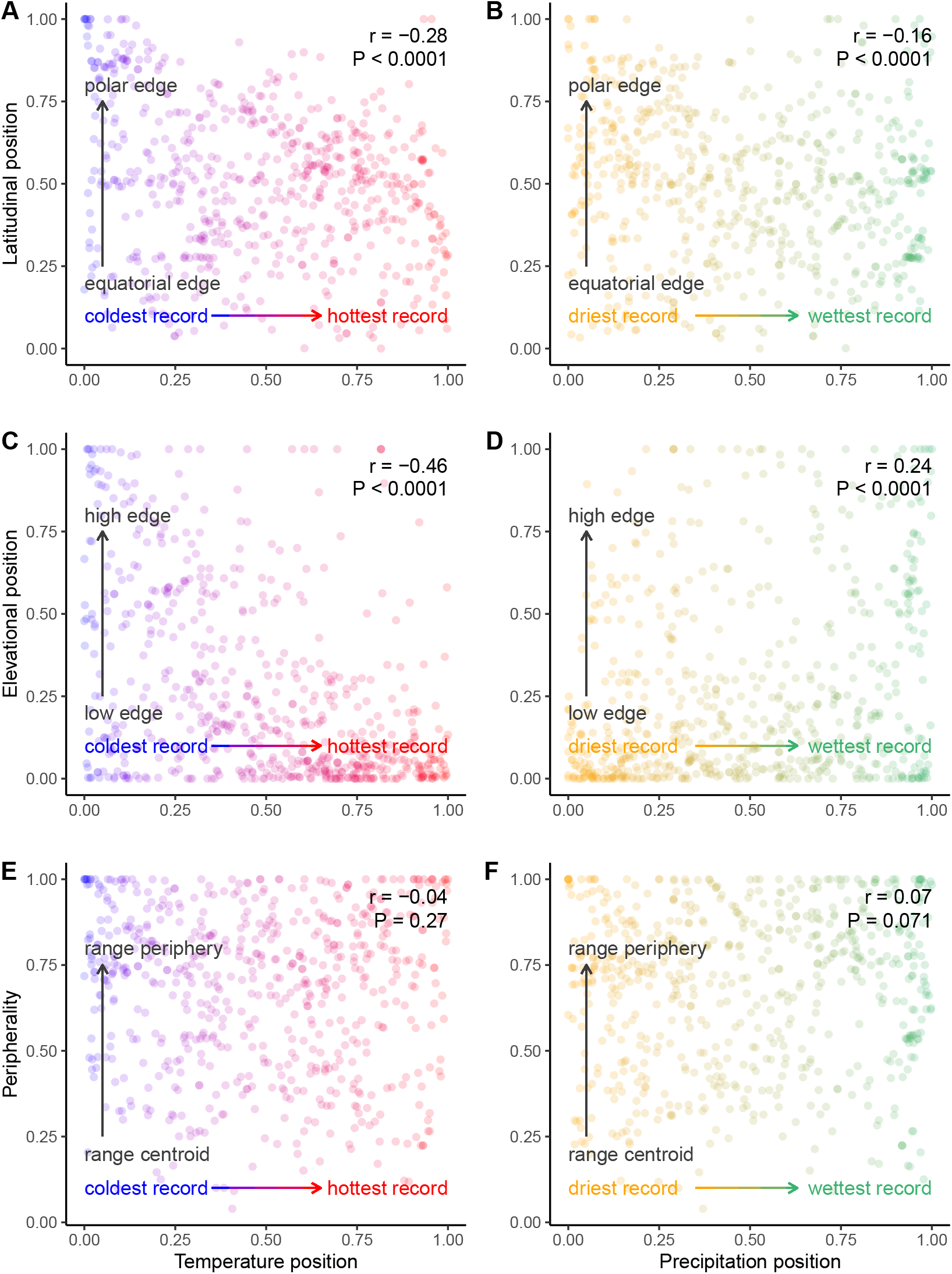
Correlation scatterplots showing weak association of spatial range position and climatic marginality. Points are transplant sites characterized by where they fall relative to occurrence records of a given species. Color coding depicts gradients from low to high values within the empirical cumulative probability distributions of mean annual temperature (red-blue gradients in panels A, C, E) or mean annual precipitation (orange-green gradients in panels B, D, F).

## Discussion

We tested how local adaptation, site quality, and population quality change across species’ ranges by harnessing two large databases: one of field transplant experiments that reveal how fitness components differ among population sources and sites, and the other of occurrence records that allowed us to categorize each transplant source and site by their positions within the species’ geographic ranges and climatic niches. We observed consistent declines in local adaptation and population quality towards high elevation and high latitude edges (Fig. 3), yet latitudinal and elevational range edges are not necessarily climatically marginal (Fig. 5). This supports the idea that legacies of range expansion can leave lasting imprints that affect current performance. Climatically marginal sites, whether they are found in the range interior or at spatial edges, have lower average performance (i.e., are of low site quality, as for cold and dry extremes) and often harbor populations whose performance is highly locally adapted (as for temperature; Fig. 4). Reinforcing the idea that range edges are not all alike (Hampe and Petit, 2005; Angert et al., 2020; Oldfather et al., 2020), peripherality itself–without consideration of direction or environment–did not strongly predict local adaptation or site quality (Fig. 3). Below we interpret the findings in more detail and touch on the implications of these results for the conservation of peripheral populations.

### Latitudinal patterns and expansion dynamics

Range edges held at equilibrium in challenging environments by the combination of selection, gene flow, and drift (Polechová, 2018) should result in edge populations being less locally adapted and of lower site and population quality (Hypothesis 1, Fig. 1). Across large spatial scales, the selection gradients that drive local adaptation are often assumed to be climatic (Gaston, 2009; Oldfather et al., 2020). Thus, if declines in local adaptation and quality at range edges are driven by marginal climates, then (a) range edges generally should be climatically marginal, and (b) spatial position should not have high explanatory power after accounting for this climatic marginality. Spatial variation in climate and fitness were inconsistent with this hypothesis; climate and range position were not tightly coupled (Fig. 5), and we detected declines in local adaptation towards poleward range edges that were not explained by climate (Fig. 3A). These results suggest either that selective forces at these edges are largely non-climatic (e.g., biotic or edaphic), or that the dynamics of spatial spread have left lasting impacts on fitness. In the latter case, range expansions should create gradients in local adaptation and population quality that reflect decreased time to adapt and increased genetic load from refugia towards recently colonized areas (Willi et al., 2018; Sheth et al., 2020), without the necessary decrease in site quality that would be expected if edges were biotically or edaphically stressful. Though post-glacial recolonization pathways and refugia can be complex (Shafer et al., 2010; Hewitt, 2011; Roberts and Hamann, 2015), for many northern hemisphere species latitude is a reasonable proxy for the timing and direction of range expansion. Consistent with the hypothesis that non-equilibrial dynamics have caused spatial variation in local adaptation and population quality at range edges (Hypothesis 2), we find that (a) local adaptation differed at equatorial versus poleward edges and (b) elevational and latitudinal position had explanatory power independent of climatic marginality (Fig. 3).

Sites near poleward range edges appeared to be similar in quality to those in other parts of the range (Fig. 3D), but populations originating from these edges performed poorly on average and were not better adapted to their home sites than foreign populations (Fig. 3A, G). These patterns are consistent with poleward-edge populations harboring greater genetic load. Genetic load has many possible sources, including immigration of maladaptive alleles (‘migration load’), genetic drift (‘drift load’), and allele surfing of deleterious mutations during spread (‘expansion load’) (Whitlock and Davis, 2011; Peischl et al., 2013). While our results cannot distinguish among these sources, accumulating evidence from genomics, field transplants, and experimental crosses suggests that range-edge populations might be particularly likely to harbor deleterious mutations and benefit from immigration (reviewed in Angert et al., 2020). This suggests that high genetic load at range edges is more likely due to expansion and drift than to migration. In contrast to poleward edges, populations from equatorial edges had an advantage over foreign populations (Fig. 3A), suggesting that they are adapted to unique aspects of their environments (although we note that we did not detect the declines in site quality that we predicted if these edges pose unique challenges). However, equatorial edge populations performed poorly on average across sites, suggesting that perhaps strong selection has reduced population sizes or that responding to local selection has come at a cost to fitness when transplanted elsewhere. Greater local adaptation at poleward edges is consistent with populations having had greater time, genetic variation, and/or spatial isolation to adapt at ‘old’ edges (de Lafontaine et al., 2018). Consistent with this interpretation, absolute latitude was also negatively correlated with local adaptation (Tables S3). Latitudinal range position maintained its explanatory power, even after accounting for the effects of absolute latitude, which may be due to different dispersal rates among regionally co-occurring species that are expanding from the same ‘glacial starting line’ (Svenning and Skov, 2004).

### Comparison of elevational and latitudinal ranges

Theoretical models for range evolution are agnostic with respect to spatial scale (e.g., whether the range dimension in question is latitudinal vs. geographic vs. elevational), and many researchers have used elevation gradients to test range limit theory (Sexton et al., 2011; Halbritter et al., 2015; Sexton et al., 2016). On the other hand, elevational and latitudinal ranges have been demonstrated to differ in connectivity and in the steepness of environmental gradients relative to species’ dispersal capacity (Hargreaves et al., 2014; Halbritter et al., 2015; Bachmann et al., 2020). Such differences lead to the prediction that elevational range edges have greater potential for gene flow to swamp divergent selection, reducing local adaptation (Bridle et al., 2009; Halbritter et al., 2015; Bachmann et al., 2020, but see Gauzere et al., 2020 and references therein regarding the prevalence of microgeographic adaptation despite gene flow). In contrast, dispersal is expected to be more limited across the larger scales associated with latitudinal ranges, possibly increasing the pervasiveness of founder effects and drift-induced genetic load. In our results, patterns of local adaptation were largely similar between latitude and elevation (Fig 3A, B). However, while populations from equatorial and polar edges did poorly across most sites (i.e., were of low population quality), population quality did not vary strongly across elevation (Fig. 3G, H). This is consistent with populations at elevational edges being less specialized and/or less subject to genetic load than those at latitudinal edges, either of which would be fostered by greater connectivity among sites across elevations compared to latitudes.

### Climate synthesis and decoupling of climate and space

Patterns of variation in site quality suggest that sites that are cool or dry relative to a species’ niche are challenging (Fig. 4C, D). Populations from warm and cool edges strongly outperformed foreign populations moved to these thermally extreme sites (Fig. 4A). Thus, while we did not find support for the hypothesis that populations at spatial range edges are strongly locally adapted (Hypothesis 3), we find an analogous pattern playing out at thermal edges. Adaptation to relatively extreme temperatures may arise if these sites consistently experience thermally extreme conditions. In contrast to responses to temperature, local adaptation did not vary with degree of precipitation marginality (Fig. 4B). This is consistent with the hypothesis that spatial and temporal variation in precipitation is less predictable (Boer and Lambert, 2008), selecting for greater plasticity instead of local adaptation (Volis et al., 2002). Alternatively, perhaps precipitation varies on a finer spatial scale than we could accurately capture at the resolution of interpolated climate grids.

### Caveats

We combined data from studies using disparate designs, encompassing different species and geographic regions. Studies estimated fitness with different components, often encompassing only some stages of the life cycle for only one or a few years. The full magnitude of local adaptation might only be apparent in lifetime fitness (Hargreaves and Eckert, 2019) and it might vary across years (Geber and Eckhart, 2005). Although these both likely added noise to our results, the former also could have introduced bias if life histories vary systematically across spatial or climatic gradients. For example, our comparisons might be more prone to underestimating local adaptation in places where life histories are slower, as is the case at high latitudes, high elevations, or low temperatures (Montesinos-Navarro et al., 2011; Laiolo and Obeso, 2017). However, in contrast to this possibility, we found high local adaptation in cold sites (Fig. 4A). Comparing local populations to foreign populations is only one component of local adaptation; another relevant comparison is the performance of populations at home to their performance away from home. Our expectations for geographic variation in local adaptation may differ slightly under these two metrics, and their comparison may yield insights into asymmetry in local fitness advantages. In our dataset, there were only half as many home-away contrasts as local-foreign, and perhaps due to reduced power we did not detect variation in local adaptation across range position metrics under this calculation (results not shown). A further caveat of this study is that spatial and taxonomic biases in our dataset were substantial, thus our conclusions are most generalizable to short-lived, small-statured plants in the northern hemisphere. More mobile or longer-lived species, which experience greater spatial and temporal environmental variation within their lifetimes, might show less local adaptation and greater phenotypic plasticity. The north-temperate bias likely increases the probability of sampling ranges still markedly influenced by post-glacial recolonization, and our results may not reflect global patterns (although glacial-interglacial cycles dramatically influenced climate and ranges even in tropical regions, Carnaval and Bates, 2007).

### Conservation considerations: climate adaptation and genetic load

Geographic peripherality has been considered a proxy for the conservation value of populations, because range edge populations are thought to occupy unique environments and as a result, may harbor unique adaptations (Lesica and Allendorf, 1995; Hardie and Hutchings, 2010). However, it has also been recognized that different types of geographic range edges may be of value for different conservation goals. Populations at leading (often poleward or high elevation) edges, for example, may be best positioned spatially to initiate warming-induced range shifts (Gibson et al., 2009), while rear edge (often equatorial) populations may harbor high genetic diversity and alleles pre-adapted to warming conditions (Hampe and Petit, 2005). There is also evidence that some range edge populations suffer from genetic load that may reduce their potential for population growth or adaptation (Willi et al., 2018). Finally, in some systems putative adaptive variation correlates more strongly with environmental than spatial predictors (Dauphin et al., 2020). As a result, developing conservation priorities for populations of any particular species requires identifying indicators of important types of diversity and balancing these with the potential costs of load in range edge populations.

Our results indicate that populations in thermally extreme sites are often strongly adapted to local conditions, but that these populations do not systematically reside at geographic range margins. Therefore, when the goal is to conserve unique climatic adaptations, it will be most effective to identify populations in climatically extreme environments, regardless of where they fall within the geographic range. This is consistent with the idea that range edges are not systematically climatically marginal due to the spatial complexity of microhabitat and landscape features (Oldfather et al., 2020). While we have only demonstrated this pattern for mean annual temperature and precipitation, it is possible that the same is true for other important climatic and non-climatic environmental pressures: the environment itself may be the best indicator of unique adaptations. In some cases, environmental extremes may coincide with range peripheries, in other cases they may not.

Our results show that poleward and equatorial edges do indeed differ in the extent to which they have adapted to their local environments. As a result, the conservation value and best conservation strategy may differ between these two types of edges. Equatorial edge populations were strongly locally adapted, consistent with the idea that these populations are reservoirs of unique genetic diversity that should be prioritized for conservation (Hampe and Petit, 2005). In contrast, polar edge populations tended to be less locally adapted and of relatively low quality. Although these patterns suggest that polar-edge populations bear relatively high genetic load, these edge populations may be important in initiating poleward range shifts forced by climate change (Gibson et al., 2009). Conservation approaches that facilitate gene flow among poleward populations could mask genetic load and facilitate future range expansions (Sexton et al., 2011; Hargreaves and Eckert, 2019).

### Future directions

Our quantitative synthesis reveals that diminished local adaptation and lower population quality (increased genetic load) may be prevalent at poleward range edges. It remains an open question whether this lower population quality compromises absolute fitness and population persistence near these range edges, and how far into the core of the range substantial load persists. Coupling estimates of load with measurement of population growth rates in natural populations or reciprocal transplants could shed light on these questions, as could experimental evolution across range microcosms. Combining experimental crosses, genomic assays, and field trials could help us understand the contributions of drift load, expansion load, and migration load at poleward edges; the relative importance of these mechanisms may determine how populations may respond to rapid environmental change or altered population connectivity. Given the well-documented prevalence of macroclimatic gradients at continental scales, the observed decoupling of climate and geographic range position is surprising. Future work should evaluate whether spatial edges are marginal along other niche axes, and whether adaptation to these factors is limited at spatial edges. Populations from thermally extreme sites strongly outperformed foreign populations, and in future it would be interesting to know if this home-site advantage comes at a cost to performance under other thermal conditions. Finally, the contrasting patterns of local adaptation to precipitation and temperature suggest that it may be interesting to measure patterns of phenotypic selection on traits involved in adaptation to different climate axes. These research avenues will further our understanding of how divergent selection and spatial evolution drive adaptation across geographic ranges, and how variation in adaptive capacity feeds back to shape species’ distributions.

## Acknowledgements

We would like to thank all of the authors of transplant studies, especially those who published their data or who provided it to us if it was not published in the form we required.

## Data accessibility

All data and code will be archived with appropriate metadata on Dryad or an equivalent public repository upon manuscript acceptance. Data and code can be provided to reviewers upon request.

