## Supplemental material for "Adaptation across geographic ranges is consistent with strong selection in marginal climates and legacies of range expansion"

### Supplementary material for: Variation in adaptation across geographic ranges is consistent with strong selection in marginal climates and legacies of range expansion

Megan Bontrager[1,6,7], Takuji Usui[1], Julie A. Lee-Yaw[2], Daniel N. Anstett[1], Haley A. Branch[1], Anna L. Hargreaves[3], Christopher D. Muir[4], and Amy L. Angert[5,7]

[1]Department of Botany and Biodiversity Research Centre, University of British Columbia, Vancouver, Canada.
[2]Department of Biological Sciences, University of Lethbridge, Lethbridge, Canada.
[3]Department of Biology, McGill University, Montreal, Canada
[4]School of Life Sciences, University of Hawaii, Honolulu, Hawaii, United States.
[5]Departments of Botany and Zoology and the Biodiversity Research Centre, University of British Columbia, Vancouver, Canada.
[6]Current affiliation: Department of Ecology and Evolutionary Biology, University of Toronto, Toronto, Canada.

March 2, 2021

#### Supplementary methods

##### Collection of fitness data and designation of local vs. foreign

When multiple fitness metrics were presented for a given species, we collected data from one representative measure in each of five categories (germination, germination and survival combined, survival, reproduction, or a composite fitness metric that incorporates at least survival and reproduction, such as population growth rate or lifetime fitness). Reproductive estimates that took into account mortality or failure to reproduce (i.e., population means that included zeros for non-reproductive or dead plants) were considered composite fitness estimates. When multiple measurements could be used for a single fitness metric (i.e., both flower counts and total seed weight were reported and could be called reproduction) we selected the one that seemed most representative of fitness, at the discretion of the data collector. If survival was reported multiple times for the same cohort (i.e., first season survival and second season survival for a perennial plant) we recorded only final survival as a proportion of the starting sample size. Germination from a single planting assessed at several time points was treated the same way. If multiple reproduction estimates were reported for a single cohort (i.e., first season fruit counts and second season fruit counts), we calculated cumulative reproduction. We calculated missing cumulative fitness metrics when components were available: we calculated a combined germination and survival estimate by multiplying those two components, and we calculated a composite metric by multiplying survival and reproduction (and germination, when available). If transplants were temporally replicated, we averaged fitness data across replicates. When studies replicated a transplant in multiple treatments (i.e., water addition or herbivore exclusion) we collected data from the treatment that most closely represented natural conditions.

Within species, we designated the local source population based on either author assignments or geographic proximity. All other populations transplanted to that site were considered foreign. Typically, this assignment was obvious–a population was collected at or very near the test site. In some studies (46 of 118) the authors did not explicitly designate a local population, so we used geographic or elevational proximity to assign one. We did not assign any population from more than 50 km of geographic distance or 100 m elevation difference from a site as local to that site. We took into account the scale at which the range was sampled when evaluating whether a suitable local assignment was available; for example, a population 10 km away from the test site would not be considered an appropriate local population if the other populations in the experiment were only moved 15 km, but might be considered an appropriate local if the other populations sampled were from hundreds of kilometers away. Even when the population assigned as local originated some distance away from the transplant site, foreign populations came from an average of 275x farther (range 1.7-5206x). Populations assigned as local were from elevations no more distant than 10% of the elevational range (average 2%) and latitudes no more distant than 3% of the latitudinal range (average 0.4%)

#### Locality cleaning methods

Prior to downloading localities, the GBIF taxonomic identifier(s) associated with each species were cross-checked with author descriptions in the transplant papers. We discarded localities that were flagged as having geospatial issues in the GBIF database, on a continent where each species is not native, or duplicates in latitude, longitude, and elevation. We also filtered out localities that were highly discontinuous with the transplant area (e.g., localities separated by major oceans from any of the transplant sites or sources). For example, if a transplant study was performed in the Americas, we filtered out Eurasian, African, and Oceanian localities. We did this because a number of species in our dataset exhibit a circumboreal distribution, but we decided the relevant geographic range for each species was the landmass contiguous with the location of the transplant sites. In one study (Vergeer and Kunin, 2013, *Arabidopsis lyrata* ssp. *petraea*), sites and sources from Iceland, the UK, and continental Europe were included, and GBIF localities from each of these landmasses were also included. We checked the locality data that we gathered against independent range maps (i.e., from field guides) or descriptions (from field guides or other published studies), and removed geographic outlier localities that might be the result of inaccurate coordinates, cultivated specimens, or misidentified specimens. GBIF records that reported an elevation of 0 were checked in Google Earth and omitted as errors if unsubstantiated. When comparing maximum and minimum elevations from GBIF with those from the literature (from field guides or other published studies), we removed the upper and lower 5% of elevational values represented in the GBIF data to avoid including potentially erroneous or outlier records in our estimates of elevational range.

#### Sensitivity test for calculating peripherality

To calculate an alternative estimate of range boundaries that allows for the range edge to have inward contours, as well as for the range to be comprised of disjunct regions, we first buffered occurrence records using the gBuffer command from the rgeos package (Bivand and Rundel, 2017) in R, with the 10th percentile of all pairwise distances between locality records used as the buffer size.

The intersection between the MCP and the resulting buffered-points polygon was then generated using the gIntersection function in the rgeos package. The peripherality of each site was calculated both with respect to the nearest edge of the global centroid of all polygons. In Sensitivity test 1, we compare our results using this estimation of peripherality to the MCP-based method used in our main analyses.

#### Supplementary results

##### Sensitivity tests

Table S14 shows comparisons of focal parameter estimates across all sensitivity tests. Local adaptation declined across latitude in all sensitivity analyses, except in sensitivity test 2, in which it declined towards both low and high latitude edges (Figure S1A). Local adaptation did not vary across elevation when elevational position was calculated with rank-based methods (sensitivity test 2), and when climate position was calculated relative to maximums and minimums, local adaptation was high at both low and high elevations (sensitivity test 3, Figure S1B). Local adaptation was not higher at thermal edges when thermal position was calculated relative to maximums and minimums (sensitivity test 3, Figure S2B).

Site quality was higher at elevational edges when elevation was calculated with a rank-based approach (sensitivity test 2, Figure S1E). Site quality was lower at thermal and precipitation extremes when climatic range position was calculated relative to maximums and minimums (sensitivity test 3, Figure S2B).

Population quality did not significantly decline at latitudinal edges in the sensitivity analyses, though the trend was similar across all tests (Figure S1G). In sensitivity test 3, where climatic and spatial predictors are all calculated relative to maximums and minimums, population quality declined at high elevations (sensitivity test 3, Figure S1H). Population quality did not increase with peripherality in sensitivity tests 1 and 2 (Figure S1I). Population quality was unrelated to precipitation position in sensitivity test 2 (Figure S2F).

##### Covariates

Local adaptation (Table S3) was negatively correlated with actual (absolute) latitude. Local adaptation also tended to decrease with increasing temperature anomaly between transplant year(s) and historical climate. The magnitude of local advantage increased with both geographic distance and thermal difference between source and site (but not precipitation difference).

Site quality (Tables S4-S8) was estimated by the performance of populations transplanted to a site (after relativizing performance of each population to its performance at other sites). The relative performance of a population was higher when that population was local to the focal site (5 of 5 subsets), and declined when the populations were moved from large geographic distances (2 of 5 subsets) or across large thermal differences (5 of 5 subsets). There was no effect of the precipitation difference between the focal site and the population's home site, or of the actual (absolute) latitude of the focal site.

Population quality (Tables S9-S13) was estimated by the performance of a population across all sites it was transplanted to (after relativizing performance of the focal population by the average performance in a site). Populations sometimes performed detectably better in their local sites (1 of 5 subsets). Population performance was negatively affected by geographic distance between the transplant site and their home site (5 of 5 subsets), but there was no effect of temperature or precipitation difference.

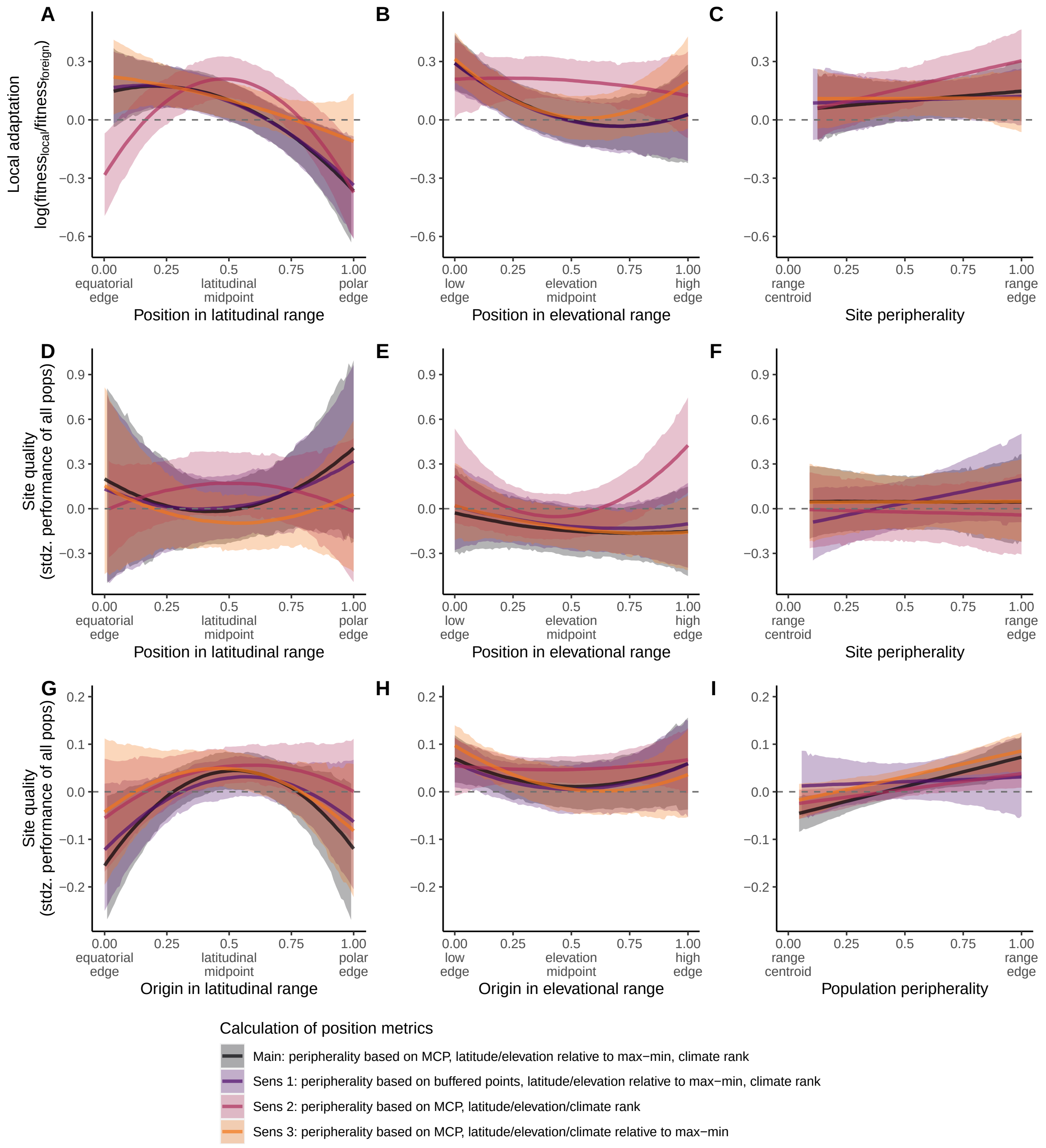


**Figure S1:** Comparison of sensitivity tests for local adaptation (A-C), site quality (D-F), and population quality (G-I) across dimensions of geographic range position defined by latitude (A, D, G), elevation (B, E, H), or a directionless peripherality metric (C, F, I). Lines and shading depict median and 95% highest density posterior interval, respectively, evaluated at mean values of the covariates. Latitudinal, elevational, thermal, and precipitation position were calculated as described in the color legend: either based on their relative position to the maximum and minimum values of occurrences or based on their rank in an empirical cumulative distribution of the values of occurrences. Peripherality was calculated relative to a minimum convex polygon (MCP) or the intersection of a MCP and buffered points. Dashed horizontal line at y=0 depicts the threshold above which populations are more fit, on average, than foreign populations at a site (A-C), than all populations in an average site (D-F), or than the average population across all sites (G-I).

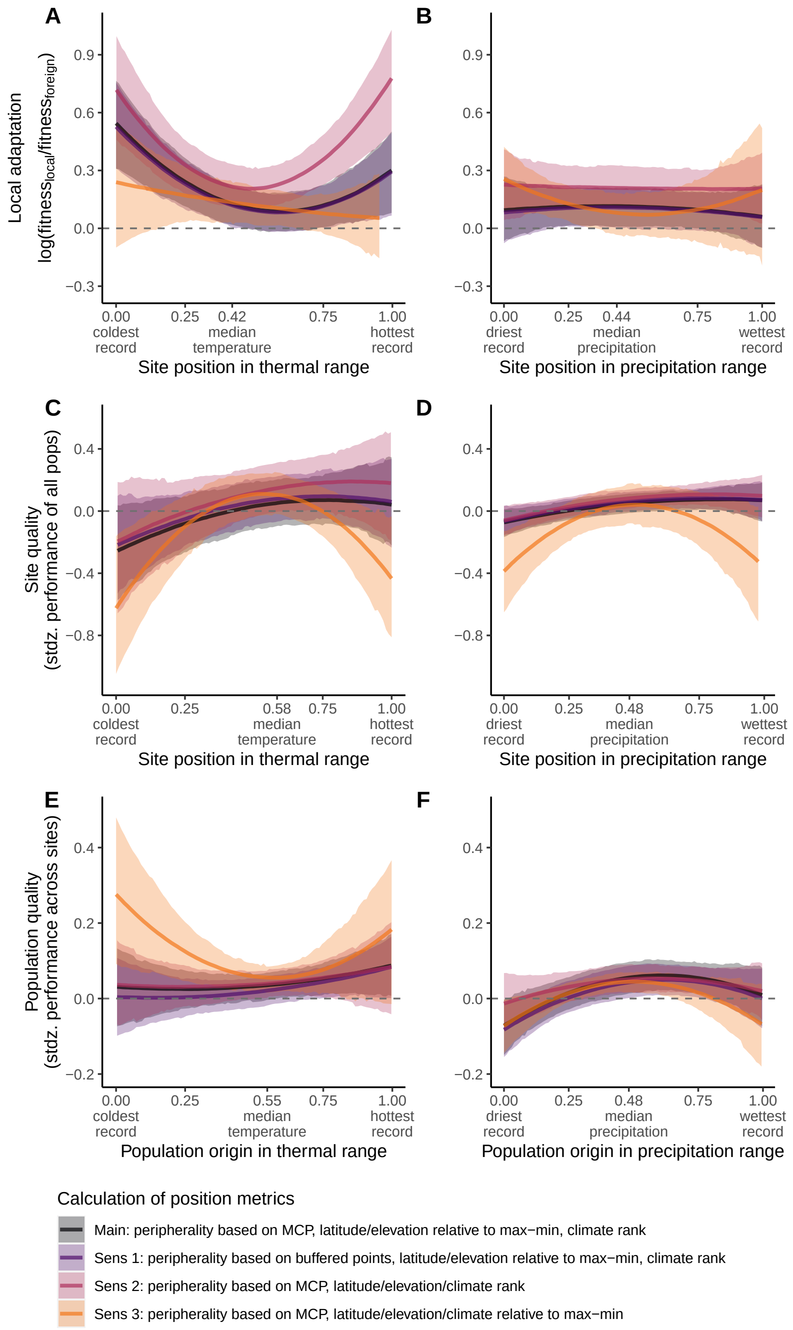


**Figure S2:** Comparison of sensitivity tests for local adaptation (A, B), site quality (C, D) and population quality (E, F) across dimensions of climatic range position defined by mean annual temperature (A, C, E) or a mean annual precipitation (B, D, F). Lines and shading depict median and 95% highest density posterior interval, respectively, evaluated at mean values of the covariates. Latitudinal, elevational, thermal, and precipitation position were calculated as described in the color legend: either based on their relative position to the maximum and minimum values of occurrences or based on their rank in an empirical cumulative distribution of the values of occurrences. Peripherality was calculated relative to a minimum convex polygon (MCP) or the intersection of a MCP and buffered points. Dashed horizontal line at y=0 depicts the threshold above which (A, B) the local population is more fit, on average, than foreign populations, or (D, E) mean fitness is greater than the average site, or (E, F) mean fitness is greater than the average population.

**Table S1:** Studies and species included in our dataset. Functional group abbreviations are: A (annual), HP (herbaceous perennial), WP (woody perennial), AR (arthropod), and F (fungus). Listed are the number of sites and sources included in the transplant experiments, the number of elevation records available to define the elevation range, the number of locality records used to define the geographic range, and the DOI of the query on GBIF.

| Study | Taxon | Functional group | No. sites | No. sources | No. elev. recs. | No. locs | GBIF DOI |
|---|---|---|---|---|---|---|---|
| Abdala-Roberts & Marquis Oecologia 2007 | Chamaecrista fasciculata | A | 3 | 3 | 53 | 1597 | 10.15468/dl.secnxl |
| Adler et al. AmJBot 2016 | Gelsemium sempervirens | HP | 2 | 9 | 111 | 400 | 10.15468/dl.37kj2g |
| Afkhami et al. EcolLett 2014 | Bromus laevipes | HP | 10 | 6 | 22 | 596 | 10.15468/dl.nt8anp |
| Agren & Schemske NewPhytol 2012 | Arabidopsis thaliana | A | 2 | 2 | 1618 | 15896 | 10.15468/dl.drv9ab |
| Alexander JBiogeography 2010 | Lactuca serriola | A | 5 | 10 | 1591 | 13998 | 10.15468/dl.hwrqxg |
| Alexander et al. JEcol 2012 | Plantago lanceolata | HP | 4 | 15 | 7120 | 27982 | 10.15468/dl.s5bn40 |
| Andersen et al. ForestEcolManag 2008 | Abies guatemalensis | WP | 2 | 9 | 182 | 254 | 10.15468/dl.kxf2rk |
| Anderson et al. AmNat 2015 | Boechera stricta | HP | 3 | 50 | 136 | 821 | 10.15468/dl.movmfs |
| Anderson et al. Evolution 2013 | Boechera stricta | HP | 2 | 2 | 136 | 821 | 10.15468/dl.movmfs |
| Angert & Schemske Evolution 2005 | Mimulus cardinalis | HP | 2 | 6 | 12 | 1001 | 10.15468/dl.4oubjy |
| Angert & Schemske Evolution 2005 | Mimulus lewisii | HP | 3 | 6 | 139 | 732 | 10.15468/dl.nddyga |
| Aparicio et al. TreeGenetGenomes 2012 | Austrocedrus chilensis | WP | 2 | 10 | 89 | 125 | 10.15468/dl.op63ym |
| Ariza & Tielborger FunctEcol 2011 | Biscutella didyma | A | 4 | 4 | 4 | 28 | 10.15468/dl.buigxr |
| Ariza & Tielborger FunctEcol 2011 | Hymenocarpos circinnatus | A | 3 | 3 | 112 | 1172 | 10.15468/dl.jm7bhp |
| Bennington et al. JEcol 2012 | Eriophorum vaginatum | HP | 6 | 6 | 153 | 869 | 10.15468/dl.topjyz |
| Bischoff et al. JEcol 2006 | Holcus lanatus | HP | 3 | 3 | 4470 | 21397 | 10.15468/dl.ce19ah |
| Bischoff et al. JEcol 2006 | Lotus corniculatus | HP | 3 | 3 | 8837 | 31599 | 10.15468/dl.xtqstk |
| Bischoff et al. JEcol 2006 | Plantago lanceolata | HP | 3 | 3 | 7120 | 27982 | 10.15468/dl.s5bn40 |
| Bischoff et al. RestorationEcol 2010 | Cichorium intybus | HP | 1 | 4 | 1278 | 12852 | 10.15468/dl.uzzfoj |
| Bischoff et al. RestorationEcol 2010 | Echium vulgare | HP | 1 | 5 | 2336 | 15368 | 10.15468/dl.i8yft0 |
| Bischoff et al. RestorationEcol 2010 | Legousia speculum-veneris | HP | 1 | 3 | 108 | 5229 | 10.15468/dl.znqs7n |
| Bischoff et al. RestorationEcol 2010 | Origanum vulgare | HP | 1 | 5 | 3127 | 13515 | 10.15468/dl.aagt28 |
| Boudry et al. JEcol 2002 | Beta vulgaris ssp. maritima | HP | 4 | 6 | 242 | 2327 | 10.15468/dl.idxmte |
| Bowman et al. JEcol 2008 | Lychnis flos-cuculi | HP | 15 | 15 | 1330 | 17587 | 10.15468/dl.sazlyc |
| Breed et al. PLoSOne 2014 | Eucalyptus gracilis | WP | 3 | 3 | 600 | 8847 | 10.15468/dl.sqaaou |
| Bresnan et al. SylvaeGenetica 1994 | Juglans nigra | WP | 7 | 66 | 44 | 1346 | 10.15468/dl.i4pauk |
| Bucharova et al. JApplEcol 2016 | Arrhenatherum elatius | HP | 4 | 7 | 3977 | 19906 | 10.15468/dl.km2tqd |
| Bucharova et al. JApplEcol 2016 | Daucus carota | HP | 4 | 8 | 2695 | 18379 | 10.15468/dl.rzf3xo |
| Bucharova et al. JApplEcol 2016 | Galium album | HP | 4 | 7 | 1123 | 11523 | 10.15468/dl.cqcurm |
| Bucharova et al. JApplEcol 2016 | Knautia arvensis | HP | 4 | 6 | 2726 | 19203 | 10.15468/dl.y4pxyu |
| Bucharova et al. JApplEcol 2016 | Lychnis flos-cuculi | HP | 4 | 7 | 1330 | 17587 | 10.15468/dl.sazlyc |
| Buckley & Bridle EcolLett 2014 | Aricia agestis | AR | 9 | 6 | 1450 | 9418 | 10.15468/dl.0kmuse |
| Byars et al. Evolution 2007 | Poa hiemata | HP | 6 | 6 | 146 | 999 | 10.15468/dl.2u06ec |
| Carter & Blair Ecosphere 2012 | Elymus canadensis | HP | 3 | 3 | 105 | 1555 | 10.15468/dl.vxj2yw |

| Study | Taxon | Functional group | No. sites | No. sources | No. elev. recs. | No. locs | GBIF DOI |
|---|---|---|---|---|---|---|---|
| Carter & Blair Ecosphere 2012 | Oligoneuron rigidum | HP | 3 | 3 | 55 | 943 | 10.15468/dl.iewffg |
| Carter & Blair Ecosphere 2012 | Sorghastrum nutans | HP | 3 | 3 | 68 | 1150 | 10.15468/dl.9pnqxd |
| Castanha et al. PlantEcolDivers 2013 | Picea engelmannii | WP | 2 | 2 | 71 | 311 | 10.15468/dl.vu6px9 |
| Castanha et al. PlantEcolDivers 2013 | Pinus flexilis | WP | 2 | 2 | 96 | 590 | 10.15468/dl.u1qts8 |
| CastellanosAcuna Ecosphere 2015 | Pinus devoniana | WP | 3 | 4 | 321 | 427 | 10.15468/dl.yetnuo |
| CastellanosAcuna Ecosphere 2015 | Pinus pseudostrobus | WP | 3 | 4 | 976 | 1252 | 10.15468/dl.mawuhu |
| Center et al. AmJBot 2016 | Quercus oleoides | WP | 2 | 4 | 441 | 588 | 10.15468/dl.yfzk6x |
| Chambers & Emery AmJBot 2016 | Vittaria appalachiana | HP | 6 | 6 | 21 | 31 | 10.15468/dl.bxlf1i |
| Chapin & Chapin Ecology 1981 | Carex aquatilis ssp. aquatilis | HP | 5 | 5 | 95 | 251 | 10.15468/dl.knwetw |
| Correia et al. AnnForestSci 2010 | Pinus pinaster | WP | 5 | 22 | 1347 | 4930 | 10.15468/dl.jm6kef |
| CostaeSilva et al. PLoSOne 2014 | Eucalyptus globulus ssp. globulus | WP | 2 | 2 | 91 | 806 | 10.15468/dl.glc2mo |
| Cremieux et al. AmJBot 2010 | Plantago lanceolata | HP | 1 | 4 | 7120 | 27982 | 10.15468/dl.s5bn40 |
| Deacon & CavenderBares PLoSOne 2015 | Quercus oleoides | WP | 2 | 2 | 441 | 588 | 10.15468/dl.yfzk6x |
| deFrenne et al. GlobalChangeBiol 2011 | Anemone nemorosa | HP | 3 | 7 | 3394 | 19015 | 10.15468/dl.iy3i5n |
| deFrenne et al. GlobalChangeBiol 2011 | Milium effusum | HP | 3 | 8 | 1669 | 17196 | 10.15468/dl.nfsjzb |
| Dunlap et al. CanJForestRes 1994 | Populus trichocarpa | WP | 2 | 8 | 81 | 1221 | 10.15468/dl.e6na78 |
| Ennos & McConnell CanJBot 1995 | Crumenulopsis sororia | F | 3 | 3 | 2 | 20 | 10.15468/dl.utshwb |
| Erfmeier & Bruelheide BiolInvasions 2010 | Rhododendron ponticum | WP | 2 | 12 | 91 | 12021 | 10.15468/dl.ansqql |
| Etterson Evolution 2004 | Chamaecrista fasciculata | A | 3 | 3 | 53 | 1597 | 10.15468/dl.secnxl |
| Fetcher et al. Biotropica 2000 | Clibadium erosum | WP | 2 | 2 | 20 | 30 | 10.15468/dl.bfxn6z |
| Fetcher et al. Biotropica 2000 | Prestoea acuminata var. montana | WP | 2 | 2 | 2 | 25 | 10.15468/dl.kujcpf |
| Fetcher et al. Biotropica 2000 | Psychotria berteriana | WP | 2 | 2 | 460 | 547 | 10.15468/dl.aq647g |
| Galen et al. Evolution 1991 | Polemonium viscosum | HP | 2 | 2 | 72 | 232 | 10.15468/dl.k2n4vk |
| Galloway & Fenster Evolution 2000 | Chamaecrista fasciculata | A | 3 | 3 | 53 | 1597 | 10.15468/dl.secnxl |
| Garrido et al. PlantEcol 2012 | Helleborus foetidus | HP | 3 | 3 | 3603 | 13055 | 10.15468/dl.7dgr71 |
| Geber & Eckhart Evolution 2005 | Clarkia xantiana ssp. parviflora | A | 2 | 2 | 0 | 41 | 10.15468/dl.slyenq |
| Geber & Eckhart Evolution 2005 | Clarkia xantiana ssp. xantiana | A | 2 | 2 | 1 | 28 | 10.15468/dl.w36eoq |
| Gibson et al. JTorreyBotSoc 2013 | Andropogon gerardii | HP | 1 | 12 | 101 | 1559 | 10.15468/dl.2uzb68 |
| Gimenez-Benavides et al. AnnBot 2007 | Silene ciliata | HP | 3 | 3 | 1159 | 1499 | 10.15468/dl.aq7dd7 |
| Gomory et al. EurJForestRes 2012 | Picea abies | WP | 4 | 11 | 4404 | 26024 | 10.15468/dl.yz6wyg |
| Gordon & Rice RestorationEcol 1998 | Aristida beyrichiana | WP | 3 | 3 | 2 | 14 | 10.15468/dl.mrgxip |
| Grassein et al. GlobalChangeBiol 2014 | Bromus erectus | HP | 2 | 2 | 3032 | 10975 | 10.15468/dl.f0bmv8 |
| Grassein et al. GlobalChangeBiol 2014 | Dactylis glomerata | HP | 2 | 2 | 13808 | 35969 | 10.15468/dl.9me17d |
| Grassein et al. GlobalChangeBiol 2014 | Festuca paniculata | HP | 2 | 2 | 1342 | 2493 | 10.15468/dl.q0vnyp |
| Grassein et al. GlobalChangeBiol 2014 | Sesleria caerulea | HP | 2 | 2 | 1592 | 9607 | 10.15468/dl.iocwal |
| Griffith & Watson JEvolBiol 2005 | Xanthium strumarium | A | 4 | 3 | 115 | 2062 | 10.15468/dl.fdflyc |
| Haggerty & Galloway JEcol 2011 | Campanulastrum americanum | A | 2 | 4 | 47 | 638 | 10.15468/dl.th8bic |
| Halbritter et al. JEvolBiol 2015 | Plantago lanceolata | HP | 3 | 9 | 7120 | 27982 | 10.15468/dl.s5bn40 |
| Halbritter et al. JEvolBiol 2015 | Plantago major | HP | 5 | 9 | 3090 | 27054 | 10.15468/dl.icoyfw |
| Hamann et al. ForestEcolManag 2000 | Alnus rubra | WP | 4 | 65 | 44 | 649 | 10.15468/dl.stulr5 |

| Study | Taxon | Functional group | No. sites | No. sources | No. elev. recs. | No. locs | GBIF DOI |
|---|---|---|---|---|---|---|---|
| Hamann et al. JEcol 2016 | Poa alpina | HP | 6 | 5 | 3551 | 9288 | 10.15468/dl.ugq4n1 |
| Hancock et al. RestorationEcol 2013 | Acacia falcata | WP | 2 | 5 | 644 | 2717 | 10.15468/dl.2ocjj8 |
| Hancock et al. RestorationEcol 2013 | Bursaria spinosa | WP | 2 | 4 | 3821 | 7524 | 10.15468/dl.pu4fcs |
| Hancock et al. RestorationEcol 2013 | Hardenbergia violacea | HP | 2 | 5 | 5500 | 7928 | 10.15468/dl.bu22xg |
| Hancock et al. RestorationEcol 2013 | Themeda australis | HP | 2 | 5 | 10 | 121 | 10.15468/dl.tbc9jr |
| Harwood et al. NewForest 1997 | Eucalyptus pellita | WP | 4 | 7 | 56 | 155 | 10.15468/dl.karlar |
| Hautier et al. JPlantEcol 2009 | Poa alpina | HP | 5 | 4 | 3551 | 9288 | 10.15468/dl.ugq4n1 |
| Hereford & Winn NewPhytol 2008 | Diodia teres | A | 6 | 6 | 107 | 366 | 10.15468/dl.3wltnl |
| Ishizuka & Goto EvolAppl 2012 | Abies sachalinensis | WP | 6 | 8 | 9 | 18 | 10.15468/dl.u9k8es |
| Jakobsson & Dinnetz EvolEcol 2005 | Carlina vulgaris | HP | 12 | 12 | 1284 | 10064 | 10.15468/dl.ne5fni |
| Jordan AmNat 1992 | Diodia teres | A | 2 | 2 | 107 | 366 | 10.15468/dl.3wltnl |
| Joshi et al. EcolLett 2001 | Dactylis glomerata | HP | 6 | 8 | 13808 | 35969 | 10.15468/dl.9me17d |
| Joshi et al. EcolLett 2001 | Plantago lanceolata | HP | 6 | 7 | 7120 | 27982 | 10.15468/dl.s5bn40 |
| Joshi et al. EcolLett 2001 | Trifolium pratense | HP | 6 | 7 | 8651 | 35488 | 10.15468/dl.h9uuj5 |
| Kim & Donohue JEcol 2013 | Erysimum capitatum | HP | 6 | 6 | 207 | 3262 | 10.15468/dl.ypxu88 |
| Koutecka & Leps Botany 2013 | Myosotis nemorosa | HP | 5 | 2 | 127 | 4905 | 10.15468/dl.mefbnj |
| Koutecka & Leps Botany 2013 | Myosotis palustris ssp. laxiflora | HP | 5 | 2 | 8 | 197 | 10.15468/dl.o2xoz9 |
| Kreyling et al. EcolEvol 2014 | Fagus sylvatica | WP | 1 | 7 | 4447 | 19819 | 10.15468/dl.za0cks |
| Lawrence & Kaye RestorationEcol 2011 | Castilleja levisecta | HP | 9 | 6 | 0 | 38 | 10.15468/dl.ehtqvg |
| Leinonen et al. AmJBot 2009 | Arabidopsis lyrata ssp. petraea | HP | 3 | 4 | 2568 | 4066 | 10.15468/dl.pftlmk |
| Liancourt & Tielborger FunctEcol 2009 | Brachypodium distachyon | A | 2 | 2 | 3845 | 9122 | 10.15468/dl.99dzbp |
| Liancourt & Tielborger FunctEcol 2009 | Bromus fasciculatus | A | 2 | 2 | 86 | 1107 | 10.15468/dl.xyx94z |
| Lopez et al. AustJBot 2007 | Pinus canariensis | WP | 4 | 21 | 275 | 2471 | 10.15468/dl.uavc9x |
| Lopez et al. PerspectPlantEcolEvolSyst 2015 | Omphalodes littoralis ssp. gallaecica | A | 5 | 5 | 3 | 32 | 10.15468/dl.wcocsb |
| Lowry et al. Evolution 2008 | Mimulus guttatus | HP | 3 | 4 | 122 | 526 | 10.15468/dl.xmvcdg |
| Lu et al. EcolEvol 2014 | Picea glauca | WP | 16 | 242 | 120 | 1123 | 10.15468/dl.cbrv7d |
| Macel et al. Ecology 2007 | Holcus lanatus | HP | 3 | 3 | 4470 | 21397 | 10.15468/dl.ce19ah |
| Macel et al. Ecology 2007 | Lotus corniculatus | HP | 3 | 3 | 8837 | 31599 | 10.15468/dl.xtqstk |
| Maes et al. PlantEcol 2014 | Milium effusum | HP | 2 | 8 | 1669 | 17196 | 10.15468/dl.nfsjzb |
| Martin & Husband Evolution 2013 | Chamerion angustifolium | HP | 9 | 11 | 97 | 1443 | 10.15468/dl.8fwdg7 |
| McCarragher et al. PhysGeog 2011 | Acer saccharum | WP | 2 | 3 | 47 | 2970 | 10.15468/dl.qdqrla |
| McGraw et al. GlobalChangeBiol 2015 | Eriophorum vaginatum | HP | 6 | 6 | 153 | 869 | 10.15468/dl.topjyz |
| McLane & Aitken EcolAppl 2012 | Pinus albicaulis | WP | 4 | 6 | 108 | 575 | 10.15468/dl.hjqqr2 |
| Milla et al. AnnBot 2009 | Lupinus angustifolius | A | 1 | 3 | 824 | 3840 | 10.15468/dl.futx81 |
| Montalvo & Ellstrand ConservBiol 2000 | Lotus scoparius | WP | 2 | 12 | 6 | 416 | 10.15468/dl.y2ps5z |
| Mylecraine et al. ForestEcolManag 2005 | Chamaecyparis thyoides | WP | 3 | 34 | 31 | 275 | 10.15468/dl.yknq1w |
| Nagamitsu et al. TreeGenetGenomes 2015 | Pinus densiflora | WP | 2 | 2 | 55 | 438 | 10.15468/dl.ur4762 |
| Pelini et al. PNAS 2009 | Erynnis propertius | AR | 6 | 6 | 23 | 325 | 10.15468/dl.neoxed |
| Pelini et al. PNAS 2009 | Papilio zelicaon | AR | 6 | 6 | 34 | 1446 | 10.15468/dl.slfy6o |
| Peterson et al. NewPhytol 2016 | Mimulus guttatus | HP | 1 | 11 | 122 | 526 | 10.15468/dl.xmvcdg |

| Study | Taxon | Functional group | No. sites | No. sources | No. elev. recs. | No. locs | GBIF DOI |
|---|---|---|---|---|---|---|---|
| Postma & Agren PNAS 2016 | Arabidopsis thaliana | A | 2 | 2 | 1618 | 15896 | 10.15468/dl.drv9ab |
| Putnam & Reich EcolMonograph 2017 | Acer saccharum | WP | 7 | 3 | 47 | 2970 | 10.15468/dl.qdqrla |
| Raabova et al. BasicApplEcol 2011 | Inula hirta | HP | 6 | 6 | 74 | 1167 | 10.15468/dl.ifized |
| Ramirez-Valiente et al. ForestEcolManag 2009 | Quercus suber | WP | 1 | 13 | 3414 | 9748 | 10.15468/dl.yut6dn |
| Reeves & Richards IntJPlantSci 2014 | Helianthus pumilus | HP | 1 | 24 | 83 | 215 | 10.15468/dl.yb0fci |
| Rice & Knapp RestorationEcol 2008 | Elymus glaucus | HP | 2 | 2 | 360 | 4424 | 10.15468/dl.4vtyyv |
| Rice & Knapp RestorationEcol 2008 | Nassella pulchra | HP | 2 | 2 | 23 | 1185 | 10.15468/dl.mpa3yz |
| Rice et al. ProcSympOakWoodl 1997 | Quercus douglasii | WP | 2 | 2 | 24 | 1096 | 10.15468/dl.xgqeo0 |
| Richards et al. Evolution 2016 | Senecio lautus | HP | 5 | 5 | 250 | 643 | 10.15468/dl.tis8tb |
| Richter et al. Oecologia 2012 | Pinus sylvestris | WP | 1 | 2 | 5629 | 31165 | 10.15468/dl.fjbh1h |
| Rosenblatt et al. EvolEcol 2016 | Melanoplus femurrubrum | AR | 2 | 2 | 139 | 631 | 10.15468/dl.6sqqza |
| Rysavy et al. JVegSci 2016 | Sarcopoterium spinosum | HP | 4 | 2 | 10 | 2607 | 10.15468/dl.8biaqp |
| Sambatti & Rice Evolution 2006 | Helianthus exilis | A | 4 | 4 | 27 | 102 | 10.15468/dl.u5fnr0 |
| Samis et al. Evolution 2016 | Camissoniopsis cheiranthifolia | HP | 4 | 8 | 4 | 775 | 10.15468/dl.nzfwth |
| Santelmann Ecology 1991 | Carex exilis | HP | 4 | 3 | 62 | 273 | 10.15468/dl.h9hrhf |
| Scheepens & Stocklin Oecologia 2013 | Campanula thyrsoides | HP | 1 | 10 | 21 | 266 | 10.15468/dl.4kk9ys |
| Schreiber et al. JApplEcol 2013 | Populus tremuloides | WP | 5 | 43 | 188 | 2466 | 10.15468/dl.eh9xxl |
| Sexton et al. PNAS 2011 | Mimulus laciniatus | A | 1 | 2 | 0 | 79 | 10.15468/dl.jozaqt |
| Smith et al. BiolCons 2005 | Lotus corniculatus | HP | 1 | 27 | 8837 | 31599 | 10.15468/dl.xtqstk |
| Stanton-Geddes et al. Ecology 2012 | Chamaecrista fasciculata | A | 3 | 5 | 53 | 1597 | 10.15468/dl.secnxl |
| Stanton-Geddes et al. PLoSOne 2012 | Chamaecrista fasciculata | A | 4 | 5 | 53 | 1597 | 10.15468/dl.secnxl |
| Streisfeld & Kohn JEvolBiol 2007 | Mimulus aurantiacus | WP | 2 | 6 | 12 | 2600 | 10.15468/dl.3trzu4 |
| Taibi et al. JEnvirManag 2016 | Pinus nigra ssp. salzmannii | WP | 3 | 5 | 220 | 907 | 10.15468/dl.x0fmzw |
| Torange et al. NewPhytol 2015 | Arabis alpina | HP | 2 | 7 | 1562 | 7199 | 10.15468/dl.yv1gqp |
| Travis & Grace EcolAppl 2010 | Spartina alterniflora | HP | 1 | 23 | 23 | 186 | 10.15468/dl.ytb5m1 |
| vanNiejenhuis & Parker CanJForestRes 1996 | Pinus banksiana | WP | 1 | 64 | 83 | 709 | 10.15468/dl.jzjzd9 |
| Vergeer & Kunin NewPhytol 2012 | Arabidopsis lyrata ssp. petraea | HP | 4 | 8 | 2568 | 4066 | 10.15468/dl.pftlmk |
| Verhoeven et al. Evolution 2004 | Hordeum spontaneum | A | 2 | 2 | 444 | 2956 | 10.15468/dl.ii2boh |
| Vizcaino-Palomar et al. PLoSOne 2014 | Pinus pinaster | WP | 2 | 2 | 1347 | 4930 | 10.15468/dl.jm6kef |
| Volis IsraelJPlantSci 2009 | Avena sterilis | A | 4 | 4 | 1376 | 9472 | 10.15468/dl.iae7cw |
| Volis et al. BiolJLinneanSoc 2002 | Hordeum spontaneum | A | 4 | 4 | 444 | 2956 | 10.15468/dl.ii2boh |
| Volis et al. PLoSOne 2015 | Triticum turgidum ssp. dicoccoides | A | 4 | 4 | 279 | 1015 | 10.15468/dl.egac3r |
| Walter et al. Evolution 2016 | Senecio pinnatifolius | A | 4 | 12 | 1179 | 7065 | 10.15468/dl.rjucko |
| Welk et al. PLoSOne 2014 | Carlina vulgaris | HP | 6 | 1 | 1284 | 10064 | 10.15468/dl.ne5fni |
| Welk et al. PLoSOne 2014 | Centaurea scabiosa | HP | 9 | 1 | 1838 | 12543 | 10.15468/dl.r4ktnx |
| Welk et al. PLoSOne 2014 | Centaurea stoebe | HP | 9 | 1 | 120 | 3584 | 10.15468/dl.lam1ls |
| Welk et al. PLoSOne 2014 | Dianthus carthusianorum | HP | 9 | 1 | 739 | 11242 | 10.15468/dl.u2nidi |
| Welk et al. PLoSOne 2014 | Dianthus deltoides | HP | 9 | 1 | 793 | 10125 | 10.15468/dl.pc0z6g |
| Welk et al. PLoSOne 2014 | Inula conyzae | HP | 8 | 1 | 730 | 7469 | 10.15468/dl.dt440v |
| Welk et al. PLoSOne 2014 | Inula hirta | HP | 8 | 1 | 74 | 1167 | 10.15468/dl.ifized |

| Study | Taxon | Functional group | No. sites | No. sources | No. elev. recs. | No. locs | GBIF DOI |
|---|---|---|---|---|---|---|---|
| Welk et al. PLoSOne 2014 | Koeleria macrantha | HP | 9 | 1 | 753 | 10550 | 10.15468/dl.ku8i15 |
| Welk et al. PLoSOne 2014 | Koeleria pyramidata | HP | 9 | 1 | 604 | 11215 | 10.15468/dl.v3gssq |
| Welk et al. PLoSOne 2014 | Scabiosa columbaria | HP | 7 | 1 | 2340 | 9973 | 10.15468/dl.setqg1 |
| Welk et al. PLoSOne 2014 | Silene nutans | HP | 9 | 1 | 2651 | 10749 | 10.15468/dl.dh5af4 |
| Welk et al. PLoSOne 2014 | Silene otites | HP | 9 | 1 | 248 | 2355 | 10.15468/dl.e2zogr |
| Wilczek et al. PNAS 2014 | Arabidopsis thaliana | A | 4 | 241 | 1618 | 15896 | 10.15468/dl.drv9ab |
| Young CanJBot 1996 | Iris douglasiana | HP | 3 | 3 | 11 | 542 | 10.15468/dl.bguizk |
| Young CanJBot 1996 | Iris innominata | HP | 3 | 3 | 21 | 120 | 10.15468/dl.obevhq |
| Zeiter & Stampfli JVegSci 2008 | Bromus erectus | HP | 3 | 3 | 3032 | 10975 | 10.15468/dl.f0bmv8 |
| Zhou et al. JEcol 2013 | Oryza rufipogon | HP | 1 | 22 | 39 | 3657 | 10.15468/dl.rvre8u |

**Table S2:** Composition of the full dataset, including the geographic, elevational, and climate distances populations were moved. Also listed are the continents where studies were conducted, the functional groups of organisms transplanted, and the fitness components that were measured. The total proportion of studies in (E) may sum to more than 1 because some studies reported multiple fitness types.

**A. Distance moved (km)**

| | Minimum | 1st Quartile | Median | Mean | 3rd Quartile | Maximum |
|---|---|---|---|---|---|---|
| Foreign sources | 0.1 | 107 | 324 | 667 | 807 | 6265 |
| Local sources | 0 | 0 | 0 | 5 | 1 | 49 |

**B. Elevations moved (m)**

| | Minimum | 1st Quartile | Median | Mean | 3rd Quartile | Maximum |
|---|---|---|---|---|---|---|
| Foreign sources | 0 | 50 | 139 | 250 | 333 | 3495 |
| Local sources | 0 | 0 | 0 | 10 | 7 | 100 |

**C. Source-site climate differences**

| | Minimum | 1st Quartile | Median | Mean | 3rd Quartile | Maximum |
|---|---|---|---|---|---|---|
| Temperature (deg.) | -18 | -0.8 | 0.2 | 0.3 | 1.3 | 15.6 |
| Precipitation (mm) | -2581 | -126 | 0 | -26 | 96 | 1594 |

**D. Continents**

| | Africa | Asia | Australia | Europe | North America | South America |
|---|---|---|---|---|---|---|
| N. sites/sources | 6 | 66 | 67 | 764 | 982 | 35 |
| Prop. sites/sources | 0.003 | 0.03 | 0.04 | 0.40 | 0.51 | 0.02 |

**E. Functional groups**

| | Annual | Herb. perennial | Woody perennial | Arthropod | Fungus |
|---|---|---|---|---|---|
| N obs. | 1608 | 3240 | 2604 | 83 | 9 |
| Prop. obs. | 0.21 | 0.43 | 0.35 | 0.01 | 0.001 |

**F. Fitness components**

| | Germination | Recruitment | Survival | Reproduction | Composite |
|---|---|---|---|---|---|
| N obs. | 346 | 481 | 3836 | 912 | 1993 |
| Prop. obs. | 0.04 | 0.06 | 0.51 | 0.12 | 0.26 |
| Prop. studies | 0.18 | 0.24 | 0.64 | 0.31 | 0.40 |

**Table S3:** Model estimates of the effects of range position metrics on **local adaptation**. Parameters of interest are plotted in Fig. 3ABC and 4AB. SE = standard error; Lower/Upper CI = lower/upper 95% highest posterior density interval from the model posterior; $\hat{R}$ = Gelman-Rubin convergence statistic (1 = convergence); ESS = effective sample size from the posterior.

| Term | Estimate | SE | Lower CI | Upper CI | $\hat{R}$ | ESS |
|---|---|---|---|---|---|---|
| Intercept | 0.11 | 0.05 | 0.02 | 0.22 | 1.00 | 1431 |
| Latitudinal position | -0.11 | 0.03 | -0.16 | -0.05 | 1.00 | 1485 |
| Latitudinal position$^2$ | -0.05 | 0.03 | -0.11 | 0.00 | 1.00 | 1487 |
| Elevation position | -0.15 | 0.04 | -0.23 | -0.07 | 1.00 | 1622 |
| Elevation position$^2$ | 0.04 | 0.02 | 0.00 | 0.08 | 1.00 | 1714 |
| Peripherality | 0.02 | 0.03 | -0.04 | 0.08 | 1.00 | 1333 |
| MAP position | -0.01 | 0.03 | -0.06 | 0.04 | 1.00 | 1249 |
| MAP position$^2$ | -0.01 | 0.03 | -0.07 | 0.05 | 1.00 | 1475 |
| MAT position | -0.10 | 0.03 | -0.16 | -0.03 | 1.00 | 1720 |
| MAT position$^2$ | 0.08 | 0.02 | 0.03 | 0.13 | 1.00 | 1206 |
| Precipitation difference | -0.01 | 0.01 | -0.02 | 0.00 | 1.00 | 1795 |
| Temperature difference | 0.03 | 0.01 | 0.02 | 0.04 | 1.00 | 2162 |
| Geographic distance | 0.05 | 0.01 | 0.04 | 0.06 | 1.00 | 2101 |
| Actual latitude | -0.05 | 0.02 | -0.09 | -0.01 | 1.00 | 1473 |
| Temperature anomaly | -0.07 | 0.03 | -0.13 | -0.01 | 1.00 | 1945 |
| Residual SD | 0.12 | 0.00 | 0.11 | 0.13 | 1.00 | 1963 |
| Student $t$ $\nu$ | 1.12 | 0.04 | 1.04 | 1.21 | 1.00 | 1959 |

**Table S4:** Model estimates of the effects of range position metrics on **site quality** for the subset of the data with good coverage of the **latitudinal range**. Latitudinal position parameters are indicated with bold text and plotted in Fig. 3D. SE = standard error; Lower/Upper CI = lower/upper 95% highest posterior density interval from the model posterior; $\hat{R}$ = Gelman-Rubin convergence statistic (1 = convergence); ESS = effective sample size from the posterior.

| Term | Estimate | SE | Lower CI | Upper CI | $\hat{R}$ | ESS |
|---|---|---|---|---|---|---|
| Intercept | 0.01 | 0.11 | -0.21 | 0.21 | 1.00 | 1553 |
| **Latitude** | **0.07** | **0.07** | **-0.07** | **0.22** | **1.00** | **1789** |
| **Latitude$^2$** | **0.05** | **0.05** | **-0.04** | **0.15** | **1.00** | **1650** |
| Elevation | 0.05 | 0.12 | -0.18 | 0.27 | 1.00 | 1680 |
| Elevation$^2$ | -0.05 | 0.04 | -0.13 | 0.02 | 1.00 | 1666 |
| Peripherality | -0.10 | 0.06 | -0.21 | 0.02 | 1.00 | 1818 |
| MAT | 0.12 | 0.08 | -0.03 | 0.29 | 1.00 | 1878 |
| MAT$^2$ | -0.06 | 0.07 | -0.20 | 0.07 | 1.00 | 1343 |
| MAP | 0.07 | 0.08 | -0.09 | 0.21 | 1.00 | 1927 |
| MAP$^2$ | -0.06 | 0.06 | -0.17 | 0.07 | 1.00 | 1717 |
| Actual latitude | -0.03 | 0.07 | -0.16 | 0.11 | 1.00 | 2000 |
| Temperature difference | -0.04 | 0.01 | -0.07 | -0.02 | 1.00 | 1848 |
| Precipitation difference | -0.00 | 0.01 | -0.02 | 0.01 | 1.00 | 2040 |
| Geographic distance | -0.01 | 0.01 | -0.02 | 0.01 | 1.00 | 1942 |
| Local to site | 0.14 | 0.04 | 0.07 | 0.22 | 1.00 | 1892 |
| Residual SD | 0.21 | 0.01 | 0.19 | 0.23 | 1.00 | 2001 |
| Student $t$ $\nu$ | 1.32 | 0.07 | 1.19 | 1.47 | 1.00 | 2052 |

**Table S5:** Model estimates of the effects of range position metrics on **site quality** for the subset of the data with good coverage of the **elevational range**. Elevational position parameters are indicated with bold text and plotted in Fig. 3E. SE = standard error; Lower/Upper CI = lower/upper 95% highest posterior density interval from the model posterior; $\hat{R}$ = Gelman-Rubin convergence statistic (1 = convergence); ESS = effective sample size from the posterior.

| Term | Estimate | SE | Lower CI | Upper CI | $\hat{R}$ | ESS |
|---|---|---|---|---|---|---|
| Intercept | -0.14 | 0.08 | -0.30 | 0.04 | 1.00 | 1295 |
| Latitude | -0.02 | 0.04 | -0.09 | 0.06 | 1.00 | 1647 |
| Latitude$^2$ | 0.04 | 0.04 | -0.03 | 0.10 | 1.00 | 1689 |
| **Elevation** | **-0.05** | **0.05** | **-0.15** | **0.06** | **1.00** | **1482** |
| **Elevation$^2$** | **0.02** | **0.04** | **-0.06** | **0.10** | **1.00** | **1711** |
| Peripherality | -0.07 | 0.05 | -0.17 | 0.03 | 1.00 | 1247 |
| MAT | 0.04 | 0.06 | -0.07 | 0.14 | 1.00 | 1636 |
| MAT$^2$ | -0.02 | 0.05 | -0.11 | 0.08 | 1.00 | 1655 |
| MAP | -0.00 | 0.05 | -0.10 | 0.10 | 1.00 | 1494 |
| MAP$^2$ | -0.01 | 0.05 | -0.09 | 0.08 | 1.00 | 1392 |
| Actual latitude | -0.03 | 0.03 | -0.09 | 0.04 | 1.00 | 1630 |
| Temperature difference | -0.04 | 0.01 | -0.06 | -0.03 | 1.00 | 1641 |
| Precipitation difference | -0.00 | 0.00 | -0.01 | 0.01 | 1.00 | 2087 |
| Geographic distance | -0.04 | 0.02 | -0.08 | -0.01 | 1.00 | 2116 |
| Local to site | 0.06 | 0.02 | 0.01 | 0.11 | 1.00 | 1909 |
| Residual SD | 0.13 | 0.01 | 0.12 | 0.14 | 1.00 | 1649 |
| Student $t$ $\nu$ | 1.00 | 0.05 | 0.91 | 1.11 | 1.00 | 1732 |

**Table S6:** Model estimates of the effects of range position metrics on **site quality** for the subset of the data with good coverage of **range peripherality**. The peripherality parameter is indicated with bold text and plotted in Fig. 3F. SE = standard error; Lower/Upper CI = lower/upper 95% highest posterior density interval from the model posterior; $\hat{R}$ = Gelman-Rubin convergence statistic (1 = convergence); ESS = effective sample size from the posterior.

| Term | Estimate | SE | Lower CI | Upper CI | $\hat{R}$ | ESS |
|---|---|---|---|---|---|---|
| Intercept | 0.05 | 0.10 | -0.14 | 0.23 | 1.00 | 1540 |
| Latitude | -0.04 | 0.07 | -0.17 | 0.09 | 1.00 | 1834 |
| Latitude$^2$ | 0.00 | 0.05 | -0.10 | 0.11 | 1.00 | 1822 |
| Elevation | -0.00 | 0.07 | -0.15 | 0.14 | 1.00 | 1631 |
| Elevation$^2$ | -0.01 | 0.03 | -0.07 | 0.05 | 1.00 | 1937 |
| **Peripherality** | **0.00** | **0.05** | **-0.10** | **0.11** | **1.00** | **1754** |
| MAT | 0.09 | 0.08 | -0.06 | 0.23 | 1.00 | 1952 |
| MAT$^2$ | -0.02 | 0.05 | -0.11 | 0.07 | 1.00 | 1791 |
| MAP | 0.06 | 0.04 | -0.02 | 0.14 | 1.00 | 1872 |
| MAP$^2$ | -0.08 | 0.04 | -0.16 | 0.01 | 1.00 | 1833 |
| Actual latitude | 0.00 | 0.10 | -0.22 | 0.20 | 1.00 | 1624 |
| Temperature difference | -0.02 | 0.01 | -0.03 | -0.00 | 1.00 | 1833 |
| Precipitation difference | 0.00 | 0.00 | -0.01 | 0.01 | 1.00 | 2048 |
| Geographic distance | -0.01 | 0.01 | -0.03 | 0.00 | 1.00 | 2034 |
| Local to site | 0.09 | 0.02 | 0.04 | 0.14 | 1.00 | 1910 |
| Residual SD | 0.10 | 0.00 | 0.09 | 0.10 | 1.00 | 2028 |
| Student $t$ $\nu$ | 1.11 | 0.05 | 1.02 | 1.21 | 1.00 | 1611 |

**Table S7:** Model estimates of the effects of range position metrics on **site quality** for the subset of the data with good coverage of the **temperature niche**. Thermal position parameters are indicated with bold text and plotted in Fig. 4C. SE = standard error; Lower/Upper CI = lower/upper 95% highest posterior density interval from the model posterior; $\hat{R}$ = Gelman-Rubin convergence statistic (1 = convergence); ESS = effective sample size from the posterior.

| Term | Estimate | SE | Lower CI | Upper CI | $\hat{R}$ | ESS |
|---|---|---|---|---|---|---|
| Intercept | 0.04 | 0.08 | -0.12 | 0.19 | 1.00 | 1470 |
| Latitude | 0.02 | 0.04 | -0.06 | 0.10 | 1.00 | 1755 |
| $\text{Latitude}^2$ | 0.03 | 0.04 | -0.05 | 0.10 | 1.00 | 1785 |
| Elevation | 0.00 | 0.08 | -0.17 | 0.15 | 1.00 | 1466 |
| $\text{Elevation}^2$ | -0.03 | 0.03 | -0.10 | 0.03 | 1.00 | 1712 |
| Peripherality | -0.06 | 0.05 | -0.15 | 0.03 | 1.00 | 1745 |
| **MAT** | **0.08** | **0.05** | **0.00** | **0.18** | **1.00** | **1592** |
| **$\mathbf{MAT^2}$** | **-0.05** | **0.04** | **-0.14** | **0.03** | **1.00** | **1386** |
| MAP | 0.07 | 0.04 | -0.00 | 0.15 | 1.00 | 1690 |
| $\text{MAP}^2$ | -0.03 | 0.05 | -0.12 | 0.06 | 1.00 | 1771 |
| Actual latitude | -0.01 | 0.03 | -0.07 | 0.05 | 1.00 | 1652 |
| Temperature difference | -0.03 | 0.01 | -0.04 | -0.01 | 1.00 | 2143 |
| Precipitation difference | 0.00 | 0.01 | -0.02 | 0.02 | 1.00 | 1944 |
| Geographic distance | -0.06 | 0.01 | -0.09 | -0.04 | 1.00 | 1797 |
| Local to site | 0.09 | 0.02 | 0.05 | 0.14 | 1.00 | 2006 |
| Residual SD | 0.14 | 0.01 | 0.13 | 0.15 | 1.00 | 1823 |
| Student $t$ $\nu$ | 1.09 | 0.05 | 1.00 | 1.21 | 1.00 | 1703 |

**Table S8:** Model estimates of the effects of range position metrics on **site quality** for the subset of the data with good coverage of the **precipitation niche**. Precipitation position parameters are indicated with bold text and plotted in Fig. 4D. SE = standard error; Lower/Upper CI = lower/upper 95% highest posterior density interval from the model posterior; $\hat{R}$ = Gelman-Rubin convergence statistic (1 = convergence); ESS = effective sample size from the posterior.

| Term | Estimate | SE | Lower CI | Upper CI | $\hat{R}$ | ESS |
|---|---|---|---|---|---|---|
| Intercept | 0.05 | 0.03 | -0.02 | 0.11 | 1.00 | 1416 |
| Latitude | -0.01 | 0.02 | -0.05 | 0.03 | 1.00 | 1377 |
| Latitude$^2$ | 0.02 | 0.02 | -0.01 | 0.06 | 1.00 | 1700 |
| Elevation | -0.02 | 0.03 | -0.08 | 0.05 | 1.00 | 1177 |
| Elevation$^2$ | -0.01 | 0.01 | -0.04 | 0.02 | 1.00 | 1542 |
| Peripherality | -0.00 | 0.02 | -0.04 | 0.04 | 1.00 | 1355 |
| MAT | 0.01 | 0.02 | -0.03 | 0.05 | 1.00 | 988 |
| MAT$^2$ | -0.04 | 0.02 | -0.08 | -0.01 | 1.00 | 1330 |
| **MAP** | **0.05** | **0.02** | **0.02** | **0.08** | **1.00** | **1595** |
| **MAP$^2$** | **-0.02** | **0.02** | **-0.05** | **0.02** | **1.00** | **1449** |
| Actual latitude | -0.01 | 0.02 | -0.04 | 0.03 | 1.00 | 1703 |
| Temperature difference | -0.04 | 0.01 | -0.05 | -0.03 | 1.00 | 1901 |
| Precipitation difference | -0.00 | 0.00 | -0.01 | 0.00 | 1.00 | 2043 |
| Geographic distance | -0.00 | 0.01 | -0.03 | 0.02 | 1.00 | 1993 |
| Local to site | 0.05 | 0.02 | 0.02 | 0.08 | 1.00 | 2130 |
| Residual SD | 0.09 | 0.00 | 0.08 | 0.10 | 1.00 | 1941 |
| Student $t$ $\nu$ | 0.98 | 0.05 | 0.89 | 1.07 | 1.00 | 2174 |

**Table S9:** Model estimates of the effects of range position metrics on **population quality** for the subset of the data with good coverage of the **latitudinal range**. Latitudinal position parameters are indicated with bold text and plotted in Fig. 3G. SE = standard error; Lower/Upper CI = lower/upper 95% highest posterior density interval from the model posterior; $\hat{R}$ = Gelman-Rubin convergence statistic (1 = convergence); ESS = effective sample size from the posterior.

| Term | Estimate | SE | Lower CI | Upper CI | $\hat{R}$ | ESS |
|---|---|---|---|---|---|---|
| Intercept | 0.04 | 0.02 | 0.00 | 0.08 | 1.00 | 1922 |
| **Latitude** | **-0.00** | **0.02** | **-0.03** | **0.03** | **1.00** | **1703** |
| **Latitude$^2$** | **-0.03** | **0.01** | **-0.06** | **-0.01** | **1.00** | **1767** |
| Elevation | -0.03 | 0.02 | -0.06 | 0.02 | 1.00 | 1798 |
| Elevation$^2$ | 0.01 | 0.01 | -0.00 | 0.03 | 1.00 | 1981 |
| Peripherality | 0.02 | 0.01 | -0.01 | 0.04 | 1.00 | 1909 |
| MAT | 0.02 | 0.02 | -0.01 | 0.05 | 1.00 | 1658 |
| MAT$^2$ | 0.03 | 0.01 | -0.00 | 0.05 | 1.00 | 1755 |
| MAP | 0.03 | 0.01 | -0.00 | 0.05 | 1.00 | 1905 |
| MAP$^2$ | -0.03 | 0.01 | -0.05 | -0.00 | 1.00 | 1920 |
| Actual latitude | 0.00 | 0.01 | -0.02 | 0.03 | 1.00 | 1387 |
| Temperature difference | 0.01 | 0.01 | -0.01 | 0.03 | 1.00 | 1835 |
| Precipitation difference | -0.00 | 0.02 | -0.03 | 0.03 | 1.00 | 2062 |
| Geographic distance | -0.09 | 0.01 | -0.11 | -0.07 | 1.00 | 1847 |
| Local to site | 0.07 | 0.04 | -0.01 | 0.13 | 1.00 | 2135 |
| Residual SD | 0.17 | 0.01 | 0.15 | 0.19 | 1.00 | 2017 |
| Student $t$ $\nu$ | 1.11 | 0.06 | 1.00 | 1.23 | 1.00 | 2033 |

**Table S10:** Model estimates of the effects of range position metrics on **population quality** for the subset of the data with good coverage of the **elevational range**. Elevational position parameters are indicated with bold text and plotted in Fig. 3H. SE = standard error; Lower/Upper CI = lower/upper 95% highest posterior density interval from the model posterior; $\hat{R}$ = Gelman-Rubin convergence statistic (1 = convergence); ESS = effective sample size from the posterior.

| Term | Estimate | SE | Lower CI | Upper CI | $\hat{R}$ | ESS |
|---|---|---|---|---|---|---|
| Intercept | 0.03 | 0.02 | -0.01 | 0.07 | 1.00 | 2042 |
| Latitude | -0.02 | 0.01 | -0.04 | 0.01 | 1.00 | 1778 |
| Latitude$^2$ | -0.03 | 0.01 | -0.05 | -0.01 | 1.00 | 1918 |
| **Elevation** | **-0.03** | **0.02** | **-0.07** | **0.01** | **1.00** | **1895** |
| **Elevation$^2$** | **0.01** | **0.01** | **-0.01** | **0.03** | **1.00** | **1993** |
| Peripherality | 0.01 | 0.01 | -0.01 | 0.04 | 1.00 | 1910 |
| MAT | 0.03 | 0.01 | 0.01 | 0.06 | 1.00 | 1926 |
| MAT$^2$ | 0.01 | 0.01 | -0.01 | 0.04 | 1.00 | 1838 |
| MAP | 0.03 | 0.01 | 0.01 | 0.06 | 1.00 | 1928 |
| MAP$^2$ | -0.03 | 0.01 | -0.05 | -0.00 | 1.00 | 1834 |
| Actual latitude | 0.02 | 0.01 | 0.01 | 0.04 | 1.00 | 1815 |
| Temperature difference | -0.01 | 0.01 | -0.03 | 0.01 | 1.00 | 1904 |
| Precipitation difference | -0.00 | 0.01 | -0.02 | 0.01 | 1.00 | 1967 |
| Geographic distance | -0.08 | 0.01 | -0.10 | -0.07 | 1.00 | 1946 |
| Local to site | 0.09 | 0.03 | 0.03 | 0.15 | 1.00 | 2042 |
| Residual SD | 0.16 | 0.01 | 0.15 | 0.18 | 1.00 | 1863 |
| Student $t$ $\nu$ | 1.17 | 0.06 | 1.07 | 1.30 | 1.00 | 1975 |

**Table S11:** Model estimates of the effects of range position metrics on **population quality** for the subset of the data with good coverage of **range peripherality**. Latitudinal position parameters are indicated with bold text and plotted in Fig. 3F. SE = standard error; Lower/Upper CI = lower/upper 95% highest posterior density interval from the model posterior; $\hat{R}$ = Gelman-Rubin convergence statistic (1 = convergence); ESS = effective sample size from the posterior.

| Term | Estimate | SE | Lower CI | Upper CI | $\hat{R}$ | ESS |
|---|---|---|---|---|---|---|
| Intercept | 0.02 | 0.01 | 0.00 | 0.04 | 1.00 | 1735 |
| Latitude | -0.02 | 0.01 | -0.03 | -0.00 | 1.00 | 2040 |
| Latitude$^2$ | -0.02 | 0.01 | -0.04 | -0.01 | 1.00 | 1730 |
| Elevation | -0.00 | 0.01 | -0.02 | 0.01 | 1.00 | 1607 |
| Elevation$^2$ | 0.00 | 0.00 | -0.01 | 0.01 | 1.00 | 1929 |
| **Peripherality** | **0.03** | **0.01** | **0.01** | **0.04** | **1.00** | **1801** |
| MAT | -0.01 | 0.01 | -0.03 | 0.01 | 1.00 | 1902 |
| MAT$^2$ | 0.01 | 0.01 | -0.00 | 0.02 | 1.00 | 1856 |
| MAP | 0.00 | 0.01 | -0.01 | 0.02 | 1.00 | 1843 |
| MAP$^2$ | 0.01 | 0.01 | 0.00 | 0.02 | 1.00 | 1890 |
| Actual latitude | -0.00 | 0.01 | -0.03 | 0.02 | 1.00 | 2084 |
| Temperature difference | -0.00 | 0.01 | -0.02 | 0.01 | 1.00 | 1855 |
| Precipitation difference | -0.01 | 0.00 | -0.02 | 0.00 | 1.00 | 1956 |
| Geographic distance | -0.04 | 0.01 | -0.06 | -0.03 | 1.00 | 1985 |
| Local to site | -0.00 | 0.02 | -0.04 | 0.05 | 1.00 | 1666 |
| Residual SD | 0.08 | 0.00 | 0.08 | 0.09 | 1.00 | 845 |
| Student $t$ $\nu$ | 1.12 | 0.06 | 1.00 | 1.24 | 1.00 | 2012 |

**Table S12:** Model estimates of the effects of range position metrics on **population quality** for the subset of the data with good coverage of the **temperature niche**. Thermal position parameters are indicated with bold text and plotted in Fig. 4E. SE = standard error; Lower/Upper CI = lower/upper 95% highest posterior density interval from the model posterior; $\hat{R}$ = Gelman-Rubin convergence statistic (1 = convergence); ESS = effective sample size from the posterior.

| Term | Estimate | SE | Lower CI | Upper CI | $\hat{R}$ | ESS |
|---|---|---|---|---|---|---|
| Intercept | 0.03 | 0.02 | -0.01 | 0.07 | 1.00 | 1801 |
| Latitude | -0.00 | 0.01 | -0.03 | 0.02 | 1.00 | 1720 |
| Latitude$^2$ | -0.01 | 0.01 | -0.03 | 0.01 | 1.00 | 1774 |
| Elevation | -0.01 | 0.02 | -0.06 | 0.03 | 1.00 | 2068 |
| Elevation$^2$ | 0.01 | 0.01 | -0.01 | 0.02 | 1.00 | 1980 |
| Peripherality | 0.01 | 0.01 | -0.01 | 0.04 | 1.00 | 1808 |
| **MAT** | **0.02** | **0.02** | **-0.01** | **0.05** | **1.00** | **1934** |
| **MAT$^2$** | **0.01** | **0.01** | **-0.02** | **0.04** | **1.00** | **1532** |
| MAP | 0.00 | 0.01 | -0.02 | 0.03 | 1.00 | 1943 |
| MAP$^2$ | -0.01 | 0.01 | -0.03 | 0.01 | 1.00 | 1913 |
| Actual latitude | 0.01 | 0.01 | -0.01 | 0.04 | 1.00 | 1966 |
| Temperature difference | 0.00 | 0.01 | -0.02 | 0.02 | 1.00 | 1841 |
| Precipitation difference | -0.01 | 0.01 | -0.02 | 0.01 | 1.00 | 2052 |
| Geographic distance | -0.09 | 0.01 | -0.11 | -0.07 | 1.00 | 1884 |
| Local to site | 0.07 | 0.04 | -0.02 | 0.15 | 1.00 | 1745 |
| Residual SD | 0.18 | 0.01 | 0.16 | 0.20 | 1.00 | 1913 |
| Student $t$ $\nu$ | 1.08 | 0.06 | 0.98 | 1.20 | 1.00 | 1859 |

**Table S13:** Model estimates of the effects of range position metrics on **population quality** for the subset of the data with good coverage of the **precipitation niche**. Precipitation position parameters are indicated with bold text and plotted in Fig. 4F. SE = standard error; Lower/Upper CI = lower/upper 95% highest posterior density interval from the model posterior; $\hat{R}$ = Gelman-Rubin convergence statistic (1 = convergence); ESS = effective sample size from the posterior.

| Term | Estimate | SE | Lower CI | Upper CI | $\hat{R}$ | ESS |
|---|---|---|---|---|---|---|
| Intercept | 0.05 | 0.02 | 0.01 | 0.09 | 1.00 | 2253 |
| Latitude | -0.01 | 0.01 | -0.03 | 0.02 | 1.00 | 1917 |
| Latitude$^2$ | -0.03 | 0.01 | -0.06 | -0.01 | 1.00 | 1878 |
| Elevation | -0.02 | 0.02 | -0.06 | 0.02 | 1.00 | 1957 |
| Elevation$^2$ | 0.00 | 0.01 | -0.02 | 0.02 | 1.00 | 2050 |
| Peripherality | 0.02 | 0.01 | -0.00 | 0.05 | 1.00 | 2094 |
| MAT | 0.00 | 0.02 | -0.03 | 0.03 | 1.00 | 2031 |
| MAT$^2$ | 0.03 | 0.01 | 0.00 | 0.06 | 1.00 | 1838 |
| **MAP** | **0.03** | **0.01** | **0.01** | **0.05** | **1.00** | **1961** |
| **MAP$^2$** | **-0.03** | **0.01** | **-0.06** | **-0.01** | **1.00** | **1904** |
| Actual latitude | 0.02 | 0.02 | -0.01 | 0.06 | 1.00 | 1992 |
| Temperature difference | -0.00 | 0.01 | -0.02 | 0.02 | 1.00 | 2062 |
| Precipitation difference | -0.01 | 0.01 | -0.03 | 0.02 | 1.00 | 1937 |
| Geographic distance | -0.09 | 0.01 | -0.12 | -0.07 | 1.00 | 2024 |
| Local to site | 0.04 | 0.03 | -0.02 | 0.10 | 1.00 | 2024 |
| Residual SD | 0.18 | 0.01 | 0.16 | 0.20 | 1.00 | 2024 |
| Student $t$ $\nu$ | 1.14 | 0.06 | 1.02 | 1.26 | 1.00 | 2054 |

**Table S14:** Comparisons of focal parameter estimates from our main models and three sensitivity analyses in which we calculated spatial or climatic predictors differently. In our main analyses, we calculated latitude and elevation relative to the maximum and minimum locations observed across all occurrences of a species, peripherality relative to an MCP, and thermal or precipitation positions based on the rank of a site in the empirical cumulative distribution of climate values associated with localities for a species. In sensitivity test 1, we calculated peripherality based on the intersection between the species' MCP and a polygon representing buffered locality data points. In sensitivity test 2, we calculated latitudinal and elevational positions with rank-based methods. In sensitivity test 3, we calculated climatic position based on temperature and precipitation maximums and minimums. Parameters with confidence intervals that do not include 0 are colored red if they are negative and blue if they are positive.

| Predictor | Main analyses | Sensitivity test 1 | Sensitivity test 2 | Sensitivity test 3 |
|---|---|---|---|---|
| **Local adaptation** | | | | |
| Latitude | -0.11 (-0.17, -0.05) | -0.11 (-0.16, -0.05) | -0.03 (-0.10, 0.04) | -0.08 (-0.13, -0.02) |
| Latitude$^2$ | -0.05 (-0.11, 0.01) | -0.04 (-0.09, 0.01) | -0.16 (-0.22, -0.09) | -0.01 (-0.06, 0.04) |
| Elevation | -0.15 (-0.23, -0.07) | -0.15 (-0.23, -0.06) | -0.01 (-0.07, 0.05) | -0.15 (-0.24, -0.07) |
| Elevation$^2$ | 0.04 (0.00, 0.08) | 0.04 (0.00, 0.08) | -0.01 (-0.05, 0.03) | 0.06 (0.02, 0.10) |
| Peripherality | 0.02 (-0.04, 0.07) | 0.01 (-0.05, 0.07) | 0.05 (0.00, 0.10) | 0.00 (-0.06, 0.06) |
| MAT | -0.10 (-0.17, -0.03) | -0.09 (-0.15, -0.03) | -0.04 (-0.12, 0.04) | -0.03 (-0.08, 0.02) |
| MAT$^2$ | 0.08 (0.04, 0.13) | 0.08 (0.04, 0.13) | 0.14 (0.08, 0.19) | 0 (-0.02, 0.03) |
| MAP | -0.01 (-0.06, 0.05) | 0.00 (-0.06, 0.05) | -0.01 (-0.06, 0.05) | -0.06 (-0.12, 0.02) |
| MAP$^2$ | -0.01 (-0.07, 0.05) | -0.01 (-0.07, 0.04) | 0 (-0.06, 0.06) | 0.02 (-0.01, 0.06) |
| **Site quality** | | | | |
| Latitude | 0.07 (-0.07, 0.22) | 0.06 (-0.09, 0.20) | 0.01 (-0.13, 0.19) | 0.01 (-0.10, 0.13) |
| Latitude$^2$ | 0.05 (-0.04, 0.15) | 0.04 (-0.05, 0.12) | -0.08 (-0.24, 0.09) | 0.04 (-0.05, 0.13) |
| Elevation | -0.05 (-0.15, 0.06) | -0.05 (-0.15, 0.05) | 0.03 (-0.07, 0.13) | -0.07 (-0.17, 0.03) |
| Elevation$^2$ | 0.02 (-0.06, 0.10) | 0.03 (-0.05, 0.11) | 0.11 (0.03, 0.18) | 0.02 (-0.06, 0.10) |
| Peripherality | 0.00 (-0.10, 0.11) | 0.07 (-0.03, 0.17) | -0.01 (-0.11, 0.09) | 0.00 (-0.10, 0.10) |
| MAT | 0.08 (-0.01, 0.18) | 0.08 (-0.02, 0.17) | 0.11 (-0.03, 0.23) | 0.02 (-0.07, 0.11) |
| MAT$^2$ | -0.05 (-0.14, 0.03) | -0.05 (-0.14, 0.04) | -0.05 (-0.14, 0.06) | -0.10 (-0.15, -0.04) |
| MAP | 0.05 (0.02, 0.08) | 0.04 (0.02, 0.08) | 0.05 (0.02, 0.08) | 0.07 (0.00, 0.14) |
| MAP$^2$ | -0.02 (-0.05, 0.02) | -0.02 (-0.06, 0.01) | -0.02 (-0.06, 0.01) | -0.06 (-0.11, -0.02) |
| **Population quality** | | | | |
| Latitude | 0.00 (-0.03, 0.03) | 0.01 (-0.02, 0.04) | 0.02 (-0.02, 0.06) | -0.01 (-0.05, 0.02) |
| Latitude$^2$ | -0.03 (-0.06, -0.01) | -0.02 (-0.05, 0.00) | -0.03 (-0.06, 0.01) | -0.02 (-0.04, 0.00) |
| Elevation | -0.03 (-0.07, 0.01) | -0.03 (-0.07, 0.00) | 0.00 (-0.02, 0.02) | -0.05 (-0.09, -0.02) |
| Elevation$^2$ | 0.01 (-0.01, 0.03) | 0.01 (0.00, 0.03) | 0.00 (-0.01, 0.02) | 0.02 (0.00, 0.04) |
| Peripherality | 0.03 (0.01, 0.04) | 0.00 (-0.02, 0.03) | 0.01 (0.00, 0.03) | 0.02 (0.01, 0.04) |
| MAT | 0.02 (-0.01, 0.05) | 0.03 (-0.01, 0.05) | 0.02 (-0.03, 0.06) | -0.01 (-0.04, 0.02) |
| MAT$^2$ | 0.01 (-0.02, 0.04) | 0.01 (-0.02, 0.04) | 0.01 (-0.03, 0.04) | 0.02 (0.00, 0.03) |
| MAP | 0.03 (0.01, 0.05) | 0.03 (0.01, 0.05) | 0.01 (-0.01, 0.04) | 0.03 (0.00, 0.05) |
| MAP$^2$ | -0.03 (-0.06, -0.01) | -0.03 (-0.06, 0.00) | -0.02 (-0.04, 0.01) | -0.02 (-0.04, -0.01) |
